# A heterozygous *CEBPA* mutation disrupting the bZIP domain causes MDS disease progression

**DOI:** 10.1101/2023.07.24.550349

**Authors:** Ruba Almaghrabi, Yara Alyayhawi, Peter Keane, Carl Ward, Rachel Bayley, Claudia Sargas, Pablo Menendez, George J. Murphy, Turki Sobahy, Mohammed A. Baghdadi, Arwa F Flemban, Saeed Kabrah, Ildem Akerman, Manoj Raghavan, Eva Barragan, Constanze Bonifer, Paloma Garcia

**Affiliations:** Institute of Cancer and Genomic Sciences, College of Medical and Dental Science, University of Birmingham, B15 2TT Birmingham, United Kingdom; Department of Laboratory Medicine (Haematology), Faculty of Applied Medical Sciences. Albaha University, Kingdom of Saudi Arabia, Al Bahah, Saudi Arabia; Department of Medical Laboratories Technology, College of Applied Medical Sciences, Jazan University, Jazan, Saudi Arabia; Molecular Biology Unit, Clinical Analysis Service, Hospital Universitario y Politécnico La Fe, 46026 Valencia, Spain; Josep Carreras Leukemia Research Institute, School of Medicine, University of Barcelona; RICORS-TERAV, ISCIII, Madrid, Spain; CIBER-ONC, ISCIII, Barcelona, Spain; Department of Biomedicine, School of Medicine, University of Barcelona, Barcelona 08036, Spain; Institució Catalana de Recerca i Estudis Avançats (ICREA), Barcelona, Spain; Section of Hematology and Oncology, Department of Medicine, Boston University School of Medicine, Boston, MA 02118, USA; Center for Regenerative Medicine (CReM), Boston University and Boston Medical Center, Boston, MA 02118, USA; Research Center, King Faisal Specialist Hospital and Research Center-Jeddah; Pathology Department, Faculty of Medicine, Umm Al-Qura University, Makkah, Kingdom of Saudi Arabia; Department of Laboratory Medicine, Faculty of Applied medical sciences, Umm Al-Qura University, Kingdom of Saudi Arabia; Institute of Metabolism and Systems Research, College of Medical and Dental Science, University of Birmingham, Birmingham, United Kingdom; Centro de Investigación Biomédica en Red de Cáncer (CIBERONC), 28029 Madrid, Spain

**Keywords:** *CEBPA*, myelodysplastic syndromes, Acute myeloid leukaemia, transcriptional regulation, erythrodysplasia, isogenic model for clonal evolution, induced pluripotent stem cells

## Abstract

Myelodysplastic syndrome disease (MDS) has a variable risk for progression to AML. Mutations in *CEBPA* are associated with a high risk of disease progression, but whether this mutation is causative for AML development is unclear. To answer this question, we generated patient-derived, MDS-specific iPSCs recapitulating the patient disease phenotype upon differentiation to blood, with hematopoietic progenitor cells showing erythroid and myeloid-dysplasia. Introduction of a frameshift mutation affecting the C/EBPα bZIP domain led to disease progression, with a reduction in clonogenic potential, block in granulocyte development and increased self-renewal capacity of erythroid progenitors. ATAC-seq revealed that the acquisition of this mutation reshaped the chromatin landscape at distal cis-regulatory regions, promoting changes in clonal composition as observed by single cell RNAseq. Our work identifies mutant *CEBPA* as causative for MDS disease progression, providing a new isogenic MDS experimental model for drug screening to improve diagnostic and therapeutic strategies.

**Highlights:**

- Development of isogenic iPSC model of clonal evolution of MDS
- Monoallelic disruption of CEBPA bZIP domain is causative for MDS disease progression
- Monoallelic disruption of CEBPA bZIP reshapes chromatin landscape
- Patient derived iPSCs recapitulate drug responsiveness

## INTRODUCTION

During ageing hematopoietic stem/progenitor cells (HSPCs) accumulate somatic mutations leading to clonal haematopoiesis of indeterminate potential (CHIP) and the development of blood disorders such as myelodysplastic syndromes (MDS). MDS are a heterogeneous group of clonal haematological diseases characterised by cytopenia and impaired haematopoiesis, with 30% of patients eventually progressing to acute myeloid leukaemia (AML) ^1, 2^. Next generation sequencing (NGS) studies have shed some light on the clonality of the disease, (i) showing that MDS founding clones contain mutations which persist when patients progress to AML ^3, 4^, (ii) that the majority of mutations are randomly acquired and not related to the pathogenesis of MDS ^3, 4^, and (iii) that commonly mutated genes such as *DNMT3A, TET2, ASXL1, TP53* and *SF3B1*, are important for initiating the disease ^5, 6^. Patients with high-risk MDS are often treated with DNA demethylating agents such as azacytidine, but this treatment proves ineffective in more than half of the patients (reviewed in^7^). The reasons behind treatment failure and AML progression have not been thoroughly investigated due to insufficient cell numbers that can be obtained from MDS patients and the lack of animal model systems that can recapitulate the spectrum of genetic mutations and chromosome alterations. The reprogramming of MDS cells into induced-pluripotent stem cells (iPSCs) has proven to be of great utility in identifying key disease associated genes such as those located in del(7q) ^8^; determining the order of mutations contributing to the clonal evolution of a MDS patient harbouring t(4;12), SF3B1, EZH2 and del5q mutations ^9^, and in constructing a phenotypic roadmap of the clonal evolution of MDS to AML with NRAS as a driver of disease progression ^10^.

However, to understand the precise role of other common mutations associated to disease progression leading to clonal heterogeneity, disease progression and high variability in disease phenotype requires further investigation. Of particular interest is the acquisition of mutations in the transcription factor CCAAT/enhancer binding protein alpha (C/EBP α) in the bZIP domain, which is required for DNA binding and dimerization. In *de novo* AML, 10-15% of patients carry mutations in *CEBPA* distributed across the entire gene, with the majority being frameshift insertions or deletions^11–13^. Most studies are based on mutations acquired in the N-terminal transactivation domain and bZIP C-terminal domain of the protein. In fact, the current World Health Organization (WHO) AML classification considers C-terminal*CEBP*A*^bZIP^*mutations as a distinct entity with favourable prognosis, better overall survival and lower risk of relapse ^14, 15^. However, a third of the CEBPA single mutations occurred between these two regions ^13^. How these mutations affect the bZIP domain in the context of MDS have not been explored and it is unknown whether they are causative for a high risk of MDS disease progression.

Here, we report the generation of MDS iPSC cells derived from a patient diagnosed with low risk MDS which are characterized by normal karyotype and carry *SRSF2, RUNX1* and *ASXL1* mutations. The patient then progressed to high risk MDS acquiring heterozygous disruption of the C/EBP α bZIP domain. We recapitulated the high-risk phenotype by introducing a similar heterozygous C/EBP α bZIP domain disruption through CRISPR-Cas9 mediated genome editing of low-risk iPSC cells, by introducing a frameshift mutation in the mid region of the protein (*CEBPA^mut^*). The haematological phenotype of these isogenic lines revealed a marked diserythropoiesis in the low-risk lines, which was accentuated during disease progression. Disruption of the C/EBP α bZIP domai n*C*i*E*n*BPA^mut^* HPCs led to a block in granulocytic differentiation, skewed differentiation towards the erythroid linage and acquisition of self-renewal properties of aberrant erythroblasts. These changes were accompanied by alterations in the chromatin of genes important for myeloid differentiation and AML such as *MAF, CEBPE, CELSR3* and *RUNX1* affecting their expression. Moreover, disruption of the C/EBPα bZIP domain led to clonal evolution. Our work therefore defines a causative role of bZIP domain *CEBPA* mutations as a driver of disease progression.

## RESULTS

### Generation of iPSCs from a low-risk MDS patient harbouring ASXL1, RUNX1 and SRSF2 mutations

To delineate a molecular and phenotypic roadmap to MDS progression, we selected a patient (MDS27) that progressed from low-risk to high-risk MDS and finally to AML, and for which longitudinal samples were available. This patient was diagnosed in July 2013 with a multilineage dysplasia and refractory anaemia and presented with a normal karyotype (low-risk MDS). In 2015, the number of blast cells increased to 9-17%; the patient had progressed to high-risk MDS and was treated with 5-Azacitine demethylating chemotherapy agents (5-Aza) but failed to respond.

Mutational screening from the sample at diagnosis (2013) revealed the presence of mutations in ASXL1 (Glu1102Asp), RUNX1 (Gly217fs) and SRSF2 (Pro95His), whilst after disease progression the patient acquired an additional mutation in the C/EBP α mid region domain (Gly257fs) changing the open reading frame. This mutation altered the sequence of the DNA binding domain and leading to a truncated protein with no dimerization domain (Figure 1A and Supplementary Table 1).

**Figure 1.**
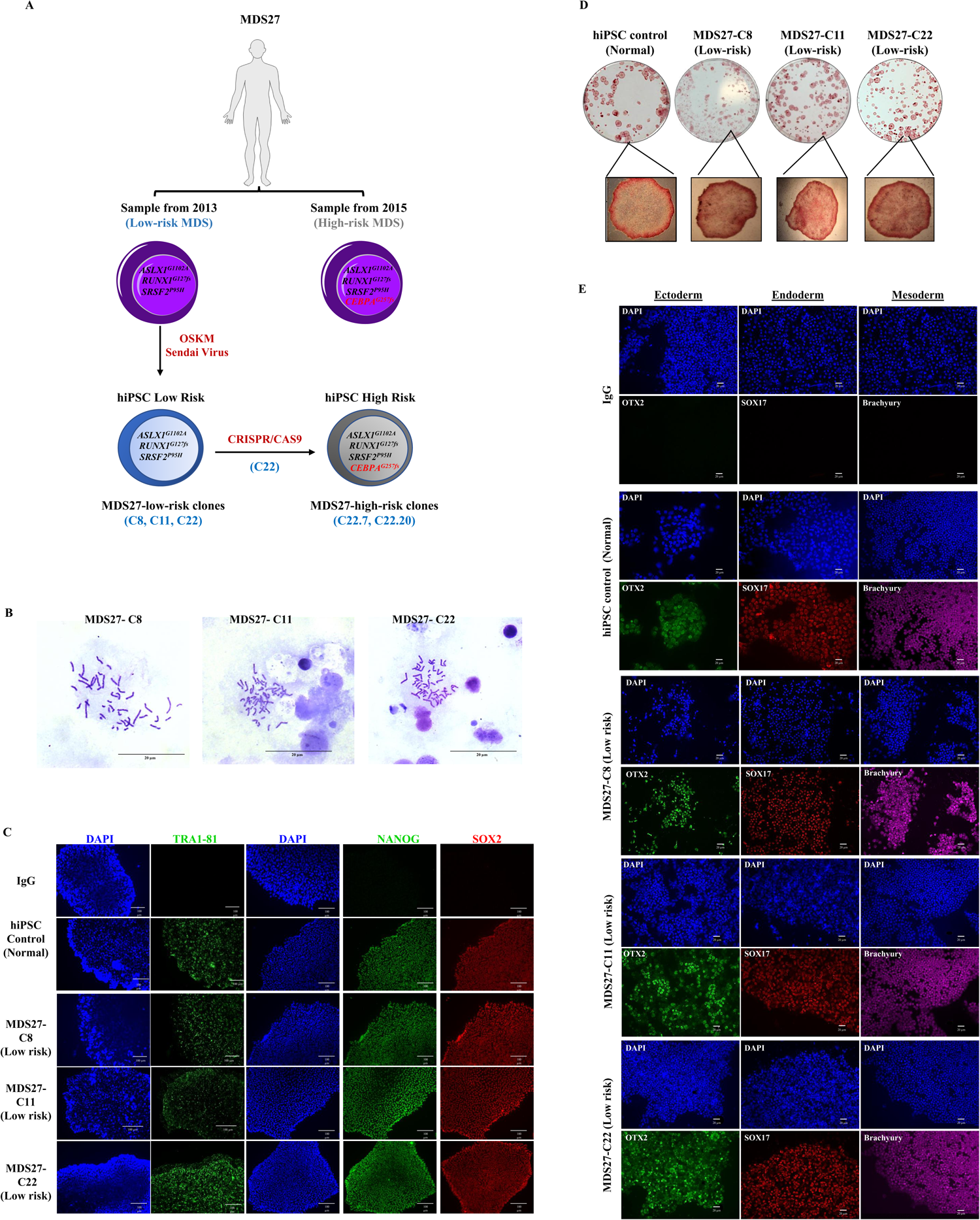
Low-risk MDS27-hiPSC lines display in vitro hallmarks of pluripotency. (A) Genotyping screening for MDS27 patient sample taken at the time of diagnoses (2013) and after disease progression (2015) as well as for iPSC clones generated from the 2013 MDS27 patient sample. (B) Chromosome spreads: Human iPSC clones generated from MDS27 showing a normal number of chromosomes. (Data represent 3 independent experiments: >25 metaphases per sample per experiment), 20 μm scale bar, 100x magnification, Leica DM6000 light microscope. (C) Representative immunofluorescence image for the staining of hiPSC control and MDS-iPSC colonies with pluripotency markers. DAPI staining is shown in blue. Scale bars, 100Cµm. 20x magnification, Leica DM6000 light microscope. N=4 independent experiments. (D) Positive expression of early pluripotent marker (alkaline phosphatase) in hiPSC control and MDS27-hiPSC lines. 10x magnification, primo vert microscope (ZEISS). N= 3 independent experiments. (E) Immunofluorescence staining of hiPSC control and MDS27-iPSC with lineage markers. The colour of SOX17 and Brachyury was modified to Red and Purple respectively using ImageJ software. 100 μm scale bar, 20x magnification, Leica DM6000. N= 4 independent experiments.

Somatic reprogramming to hiPSC was performed on peripheral blood mononuclear cells (PBMCs) obtained from patient MDS27 at the time of diagnosis (2013) using Sendai virus (SeV), a non-integrative system^16^. 24 iPSC individual colonies were isolated, expanded and established as MDS27 low-risk iPSC lines. A genotyping screening confirmed that all the hiPSC colonies examined (15 colonies in total) were positive for the same *ASXL1, RUNX1* and *SRSF2* mutations present in the original PBMCs (Supplementary Table 1). Mutational screening confirmed that no isogenic healthy control clones from MDS27 were generated nor colonies with different combinations of these mutations, despite performing somatic reprogramming on three different occasions and with two different non-integrative methods (episomal and SeV) ^17^ (Supplementary Table 1). These findings suggest that cells harbouring these three mutations either seem to be more amenable to reprogramming or, more likely, that *ASXL1, RUNX1, SRSF2* mutated cells were already a predominant clone in the patient at the time of diagnosis.

Given that no isogenic healthy iPSCs were obtained, a published non-isogenic iPSC line generated from healthy PBMCs was used as a control, namely BU3.10 (CREM003i-BU3C2; WiCell). Three different hiPSC clones generated from low-risk MDS27 by SeV (C8, C11, C22) were used for further characterisation. All three clones maintained a normal karyotype (Figure 1B), were positive for the expression of pluripotent markers TRA1-81, NANOG and SOX2 ^18, 19^ (Figure 1C) and showed positive alkaline phosphatase staining (AP) ^20, 21^ (Figure 1D). Furthermore, all clones were capable of differentiation to the three germ layers as shown by the expression of OTX2, SOX17 and Brachyury when differentiated towards ectoderm, endoderm, and mesoderm, respectively ^22–25^ (Figure 1E). These clones exhibit all aspects of characteristics of pluripotent stem cells: They are morphologically undifferentiated, positive for AP, express the pluripotent markers and can differentiate into the three germ lineages.

### Low-risk MDS27-iPSC harbouring ASXL1, RUNX1 and SRSF2 mutations display normal hematopoietic progenitor potential

To assess the differentiation potential of the MDS27-iPSC clones towards hematopoietic progenitor cells (HPCs), we used the STEMdiff Hematopoietic protocol from STEM Cell Technology (Supplementary Figure 1A). The course of differentiation was monitored using CD34 (an HSC and early progenitor marker), CD43 (an early hematopoietic marker which persists in differentiating precursor cells) ^26^, and CD45 (the key marker of human hematopoietic cells) ^27^. Expression of these markers was assessed by flow cytometric analysis on progressive days (10, 12 and 14) to determine the dynamics of hematopoietic differentiation. 3 different iPSC clones derived from the MDS27 patient sample (C8, C11 and C22) were investigated to control for possible variations due to the reprogramming process. Throughout the hematopoietic differentiation, the hiPSC control and MDS27 hiPSC clones behaved very similarly (Supplementary Figure 1B). Typically, by day 14, around 90% of cells were CD43+ and 25-35% were CD45 ^+^/CD34^+^ (Supplementary Figure 1C). In summary, this analysis demonstrates that we successfully differentiated MDS27 hiPSC clones towards early and late hematopoietic cells at similar rates as hiPSC control and thus, that *ASLX1, RUNX1* and *SRSF2* mutations do not affect the generation of HPCs from hiPSC-MDS27 clones.

We next proceeded to assess their myeloid maturation potential of iPSC generated HPCs. HSPCs from control and MDS27 hiPSC clones were obtained at day 12 and plated in methylcellulose medium enriched with recombinant cytokines that induce the differentiation into colony forming units of mixed lineage (CFU-GEMM), bursting forming units of committed erythroid (BFU-E), and colony forming units of myeloid lineage progenitors (CFU-G, CFU-M, CFU-GM) (Supplementary Figure 2A). On day 14, the number of colonies were counted and scored based on their phenotypic characteristics. All clones derived from the MDS27 patient sample displayed a similar behaviour (Figure 2A and 2B). Consistently throughout all independent experiments, we observed a lower number of colonies with HPCs derived from all three MDS27-hiPSC clones compared to the healthy hiPSC control. The reduction in the number of myeloid CFUs indicated an impaired hematopoietic colony-forming capacity of these clones (Figure 2A). All clones derived from the MDS27 patient sample were able to generate all types of CFU(Fsigure 2B), but the proportion of CFU-M and CFU-G was lower in comparison to the control line (Figure 2C). Despite the decrease in hematopoietic colony number in MDS27-hiPSC clones, no difference in the morphology and size of the colonies formed was observe dSu(pplementary Figure 2B), although we noticed an evident bias towards the formation of erythroid colonies versus myeloid colonies.

**Figure 2:**
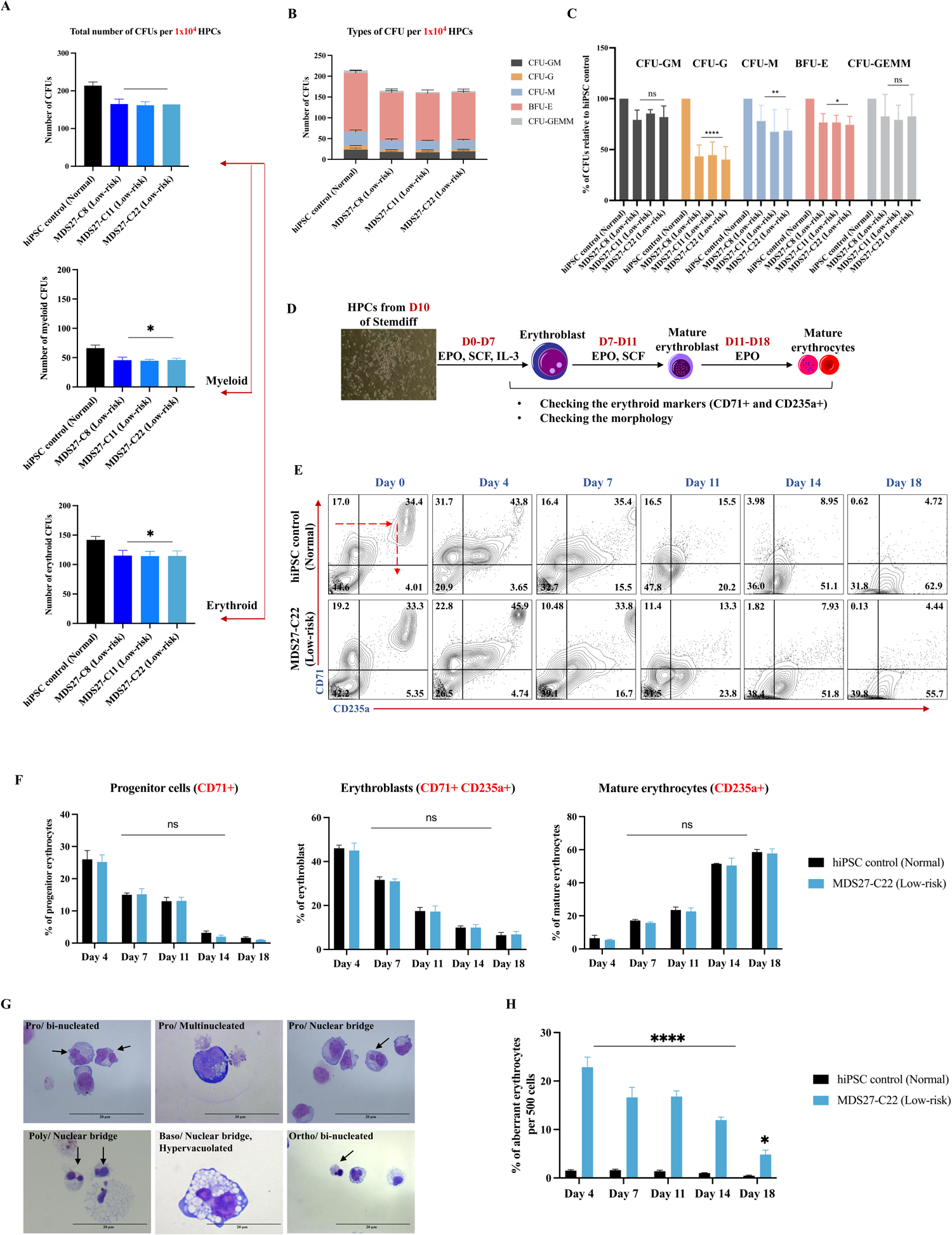
Low-risk MDS27-hiPSC lines are able to differentiation into hematopoietic progenitors in semi-solid medium and into erythroid cells in the liquid culture. (A) Total number of CFUs from 10 ^4^ –iPSC-HSPC cells grown for 14 days in semi solid medium. Statistical results are presented as mean ± SEM. * p< 0.05; ****p <0.0001 and (ns, no significant), One-way ANOVA with multiple comparisons. N= 4 independent experiments. (B) Number of each type of CFUs after 14 days in semisolid medium. Mean and SEM of different lines are shown. N= 4 independent experiments. (C) Relative percentage of each type of CFUs for 1×10 ^4^ of HSPCs after 14 days in semisolid media. Statistical results are presented as mean ± SEM and **** p < 0. 0001, **p< 0.01, * p< 0.05 and (ns, no significant), Two-way ANOVA with multiple comparisons. N= 4 independent experiments. (D) Schematic representation of the experimental set up for the erythroid differentiation of HSPCs. (E) Flow cytometry dot plots for CD71 and CD235a markers on Day 0, 4, 7, 11, 14 during erythroid differentiation. N= 4 independent experiments. (F) Percentage of erythrocyte (CD71^+^), erythroblasts (CD71 ^+^ CD235a^+^) and mature erythrocytes (CD235a^+^) during erythroid differentiation. Mean and SEM are show, (ns, no significant). N= 4 independent experiments. (G) Diff-quick stained cytospins showing common aberrant morphology (black arrow) observe in MDS27-C22. The pictures were taken by Leica DM6000 at 40x and 100x magnification, 20 μm scale bar. N= 3 independent experiments. (H) Percentage of erythroid cells with aberrant morphology. Statistical results are presented as mean ± SEM. ***p <0.0001 and ** p <0.001, Two-way ANOVA with multiple comparisons. N=4 independent experiments.

### Low risk MDS27-hiPSC harbouring ASXL1, RUNX1 and SRSF2 mutations display aberrant erythroid differentiation

Given that anaemia is one of the most common features of MDS patients, we sought to determine whether erythropoiesis might be impacted in patient-derived low-risk clones. To this end, we evaluated the erythroid differentiation potential of generated HPCs in liquid culture. As all the clones generated from low-risk MDS behaved similarly in terms of clonogenic capacity and differentiation potential, one clone was selected for further study (clone 22). Day 10 HPCs obtained from control and MDS27-C22 (low risk) hiPSC were cultured in an established 3-phase erythropoiesis liquid culture over 18 days (Figure 2D). The culture was monitored every 3-4 days, and the distribution over different erythroid maturation stages was assessed by measuring the expression of CD71 (progenitor marker) and CD235a (Glycophorin A). After culturing HPCs in erythroid media for 7 days, roughly 15% of cells were CD71 ^+^ (erythroid progenitors) and 40% of cells co-expressed CD71 ^+^ and CD235a^+^ (erythroblasts), while the expression of the mature marker (CD235a ^+^) was very low at this early stage of the differentiation. This cellular distribution was very similar for both hiPSC control and MDS27-C22 hiPSC (Figure 2E and 2F). As differentiation continued, the percentage of erythroblasts decreased with a concomitant increase in the percentage of the mature cells. On the final day of erythroid differentiation (day 18), the erythroid lineage cells became more mature, with over 55% of the cells expressing CD235a^+^ and only 5% of cells being still at the erythroblast stage (CD71^+^ CD235a ^+^) (Figure 2E and 2F). Analysis of these surface markers by flow cytometry did not highlight any immunophenotypic differences in MDS27 clones compared to the control (Figure 2F).

Morphological evaluation confirmed that both control and MDS27-C22 hiPSCs had given rise to a heterogeneous population of erythroid cells. All the stages of maturation appeared during the several days of the differentiation (Supplementary Figure 2C). Nucleated erythroblasts were observed in the cultures from day 4 in both hiPSC control and MDS27-C22 hiPSC clones. Enucleated erythroid cells (mature cells) were observed as early as day 7 (Supplementary Figure 2C, yellow arrows). Moreover, the morphological analysis revealed that the erythroid differentiation of MDS27-C22 hiPSCs was dysplastic as cells presented aberrant morphology (binucleated, multinucleated and nuclear bridges) (Figure 2G, 2H and Supplementary Figure 2C), common dysplastic features found on MDS patients ^28^. The increase in cells with aberrant morphology was evident in the erythroid cultures from MDS27-C22 iPSCs compared to control hiPSC at all time points (Day 4, 7, 11, 14 and 18) (Figure 2H). These data indicate that *SRSF2, ASLX1* and *RUNX1* mutations promote an aberrant maturation of erythroid cells in low-risk MDS, and importantly, that the disease phenotype could be reproduced *in vitro*.

### Generation of high-risk MDS27 iPSC by CRISPR-Cas9 mediated genome editing

Mutational screening of the MDS27 patient sample obtained in 2015 revealed the acquisition of a heterozygous C/EBPα mutation in the mid region of the protein disrupting the open reading frame of the bZIP domain (*CEBPA^mut^*). Somatic reprogramming to iPSC was performed on PBMCs obtained at this stage of disease using SeV, but iPSC generated were not stable in culture and suffered spontaneous non-specific differentiation. Thus, to obtain cells before and after disease progression with the same genetic background (harbouring mutations in *ASXL1, RUNX1* and *SRSF2*), and to understand the contribution of mutations disrupting the C/EBP α b Z I P domain to the disease progression, we applied CRISPR-Cas9 technology to introduce a *C E B P A* mid-region mutation disrupting the bZIP domain into the MDS27-C22 hiPSCs generated, mimicking the disruption of the bZIP domain observed in the MDS patient. 24 hours post-nucleofection, cells were dissociated and the GFP ^+^ cells sorted by FACS (Supplementary Figure 3 A and 3B). 3 clones (C22.5, C22.7 and C22.20) out of 7 were found to contain a heterozygous mutation based on the T7EI assay (Supplementary Figure 3C). Sanger sequencing confirmed that all 3 clones harboured the deletion in the target site in one allele of the *C/EBPα* gene (Supplementary Figure 3D). The sequencing data showed that MDS27-C22.5 contained multiple integrations and deletions whilst MDS27-C22.7 and MDS27-C22.20 had both a 43bp deletion in the targeted region of interest with a consequent premature termination of C/EBPα protein, altering the DNA binding domain of C/EBPα, and thus its functionality (Supplementary Figure 3E). Characterization of the new clones generated (MDS27-C22 CRISPR control, MDS27-C22.7 (High risk 1) and MDS27-C22.20 (High risk 2)) confirmed the absence of aneuploidy (Supplementary Figure 4A) and positive expression of pluripotent protein markers (TRA1-81, SOX2 and NANOG) (Supplementary Figure 4B). Additionally, differentiation of those cells towards HPCs revealed no significant differences in the expression of hematopoietic markers at day 14, indicating that inclusion of the *CEBPA* mutation in cells harbouring mutations in *ASXL1, RUNX1, SRSF2* did not affect the generation of HPCs (Supplementary Figure 4C).

**Figure 3:**
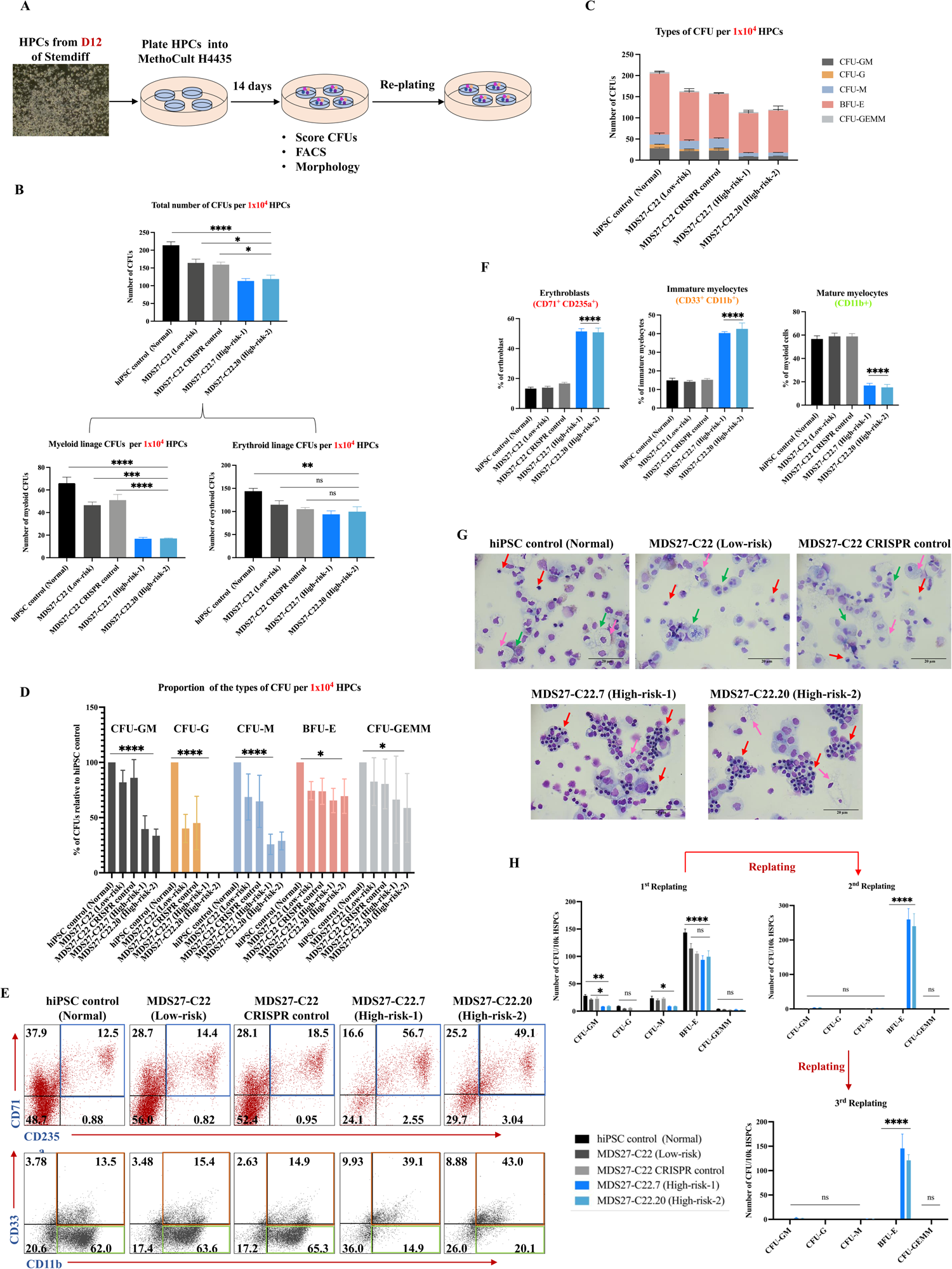
Erythroid-biased differentiation and increased self-renewal capacity of high-risk MDS-iPSC containing a *C/EBPα* mutation. (A) Schematic representation of the experimental design. (B) Total number of CFUs from 10 ^4^ –iPSC-HSPCs grown for 14 days in semi solid medium. Statistical results are presented as mean ± SEM. **** p < 0, ***p <0.0001, ** p <0.001, * p< 0.05 and (ns, no significant), One-way ANOVA. N=4 independent experiments. (C) Number of each type of CFUs after 14 days in semisolid medium. Mean and SEM of different lines are shown. Mean and SEM of different lines are shown. N= 4 independent experiments. (D) Relative percentage of each type of CFUs for 1×10 ^4^ of HSPCs after 14 days in semisolid media. Statistical results are presented as mean ± SEM and **** p < 0.0001 and * p< 0.05, Two-way ANOVA. N= 4 independent experiments. (E) Representative flow cytometry panels showing the percentage of erythroblasts cells (blue square) (CD71 ^+^CD235a^+^), immature myeloid cells, orange square (CD33 ^+^ CD11b^+^), and mature myeloid cells, green rectangle (CD11b ^+^) obtained from colony assays. N= 3 independent experiments. (F) Fraction of erythroblasts (CD71 ^+^ CD235a^+^), immature myeloid (CD3^+^3CD11b^+^) and mature myeloid cells (CD11b ^+^). Mean and SEM are shown. **** p < 0.0001, One-way ANOVA with multiple comparisons. N= 3 independent experiments. (G) Diff-quick stained cytospins from colony assay showing erythroid cells (Red arrows), granulocytes (Green arrows) and monocytes (Pink arrows). The pictures were taken Leica DM6000 at 40x, 20 μm scale bar. N= 3 independent experiments. (H) Number of CFUs obtained from control, WT clones and mutant clones after second and third re-plating, each maintained for 14 days. Mean and SEM are shown. **** p < 0.0001, ** p<0.001, * p 0.05 and (ns, no significant), Two-way ANOVA. N= 3 independent experiments.

### Erythroid-biased differentiation and increased self-renewal capacity of high-risk MDS-iPSC containing C/EBPα mutations

We next sought to define whether the C/EBPα mutation would alter the hematopoietic differentiation phenotype observed in cells containing mutations in *ASXL1*, *SRSF2* and *RUNX1*, and thus serve as a model system to study disease progression. The differentiation potential and clonogenic capacity of HPCs derived from hiPSC control, MDS27-C22 (Low risk), MDS27-C22 CRISPR control, MDS27-C22.7 (High risk-1) and MDS27-C22.20 (High risk-2) hiPSCs was assessed by a colony assay in methylcellulose medium (Figure 3A). HPCs from both controls, and low risk iPSCs, gave rise to both myeloid and erythroid CFUs whilst the HPCs derived from high risk iPSCs (containing the *CEBPA* mutation), exhibited a significant reduction in their clonogenic capacity affecting mainly the myeloid lineage (Figure 3B and 3C and Supplementary Figure 5A). Scoring the type of CFUs showed that *CEBPA* mutant clones were not able to differentiate to CFU-G but were mainly forming BFU-E (Figure 3C and 3D).

To corroborate this finding, cells were collected to assess morphology or to be stained with antibodies against the myeloid (CD33, CD11b) and erythroid markers (CD71 and CD235a) to assess their expression by flow cytometry. This analysis revealed that *CEBPA* mutant clones gave rise to a lower percentage of myeloid cells composed mostly of immature cells CD3^+^3CD11b^+^ (orange square) whilst the hiPSC control, MDS27-C22 and MDS27-C22 CRISPR control hiPSCs, were able to form mature myeloid CD11b ^+^ cell (green rectangle) (Figure 3E&F). The flow cytometry analysis confirmed that *CEBPA* mutant clones, MDS27-C22.7 and MDS27-C22.20, could generate significantly erythroid colonies in methylcellulose culture with approximately half being erythroblast cells (CD71 ^+^CD235a^+^, blue square) (Figure 3E&F). Erythroid-biased differentiation was also evident by the increase in erythroblast cells observed with diff-quick stainingFi(gure 3G).

To assess whether the *CEBPA* mutation conferred an increase in self-renewal capacity, serial re-plating in methylcellulose was performed. Secondary re-plating resulted in the generation of BFU-E at a frequency of 264±31 and 243±36 colonies in MDS27-C22.7 and MDS27-C22.20 hiPSCs respectively, whilst both controls and low risk hiPSCs, had very limited re-plating capacity with a frequency of 2 to 3 colonies of CFU-GM and no BFUE colonies (Figure 3H). In addition, a dramatic increase in the number of cells was observed in the *CEBPA* mutant lines from first re-plating to second re-plating. This result was in contrast with the other iPSC cell lines (hiPSC control, MDS27-C22 (Low risk) and MDS27-C22 CRSPR control) in which a decrease in the total number of cells was found from the first to the second re-plating (Supplementary Figure 5B). Cells from the *CEBPA* mutant clones (high risk) were able to proliferate in third-replating whereby the number of BFU-E had decreased to 149±29 and 133±18 colonies, respectively and cells lost their proliferation capacity at the fourth re-plating (Figure 3H).

To determine whether the *CEBPA* mutation alone was responsible for the gain in self-renewal capacity, a *CEBPA* mutation disrupting the bZIP domain was generated in BU3.10 wild type iPSC cells (BU3.10-*C/EBPα^mut^* C7 and C12). The BU3.10-*C/EBPα* mutant iPSC lines were able to differentiate to HPCs in equal proportions as their isogenic controls (Supplementary Figure 6A) but displayed a reduction on CFU-G and BFU-E colonies (Supplementary Figure 6B). However, in contrast to MDS27-C22.7 and MDS27-C22.20 (high risk) iPSCs, BU3.10-*C/EB P ^m^α^ut^* C7 and C12 did not form colonies after secondary and tertiary re-plating (Supplementary Figure 6C).

Taken together, these results demonstrate that a heterozygous *CEBPA* mutation which disrupts the bZIP domain affects myeloid differentiation as expected from previous studies ^11^, thus validating our model. Our results also show that introducing the *CEBPA* mutation on the background of additional mutations (*ASXL1, RUNX1, SRSF2*) but not by itself increased the self-renewal capacity of the committed progenitor cells and blocked myeloid differentiation, which is the main characteristic of high-risk MDS and AML.

### Reduced myeloid differentiation capacity of high-risk MDS-iPSC containing C/EBPα mutation

Colony assays revealed that the *C/EBPα* mutation in combination with *ASLX1, RUNX1* and *SRSF2* mutations affect the myeloid differentiation capacity of HPCs (Figure 3). Thus, to get a better understanding of how these mutations reduce myeloid differentiation potential, we performed myeloid lineage differentiation in liquid culture (Figure 4A). The emergence of myeloid cells was assessed by flow cytometric analysis throughout the differentiation time course (day 0, 4 and 7) by looking at the percentage of pure granulocytic cells (CD11b ^+^CD14^-^) and monocytic cells (CD11b ^-^ CD14^+^/CD11b^+^CD14^+^). By day 7 all hiPSC lines without the *C/EBPα* mutation were capable of differentiation to granulocytes (around 40%). In contrast, the *C/EBPα^mut^* hiPSC lines (high risk) had a significantly limited capacity to produce granulocytes and their differentiation seemed to be skewed to the production of monocytes (Figure 4B, 4C).

**Figure 4:**
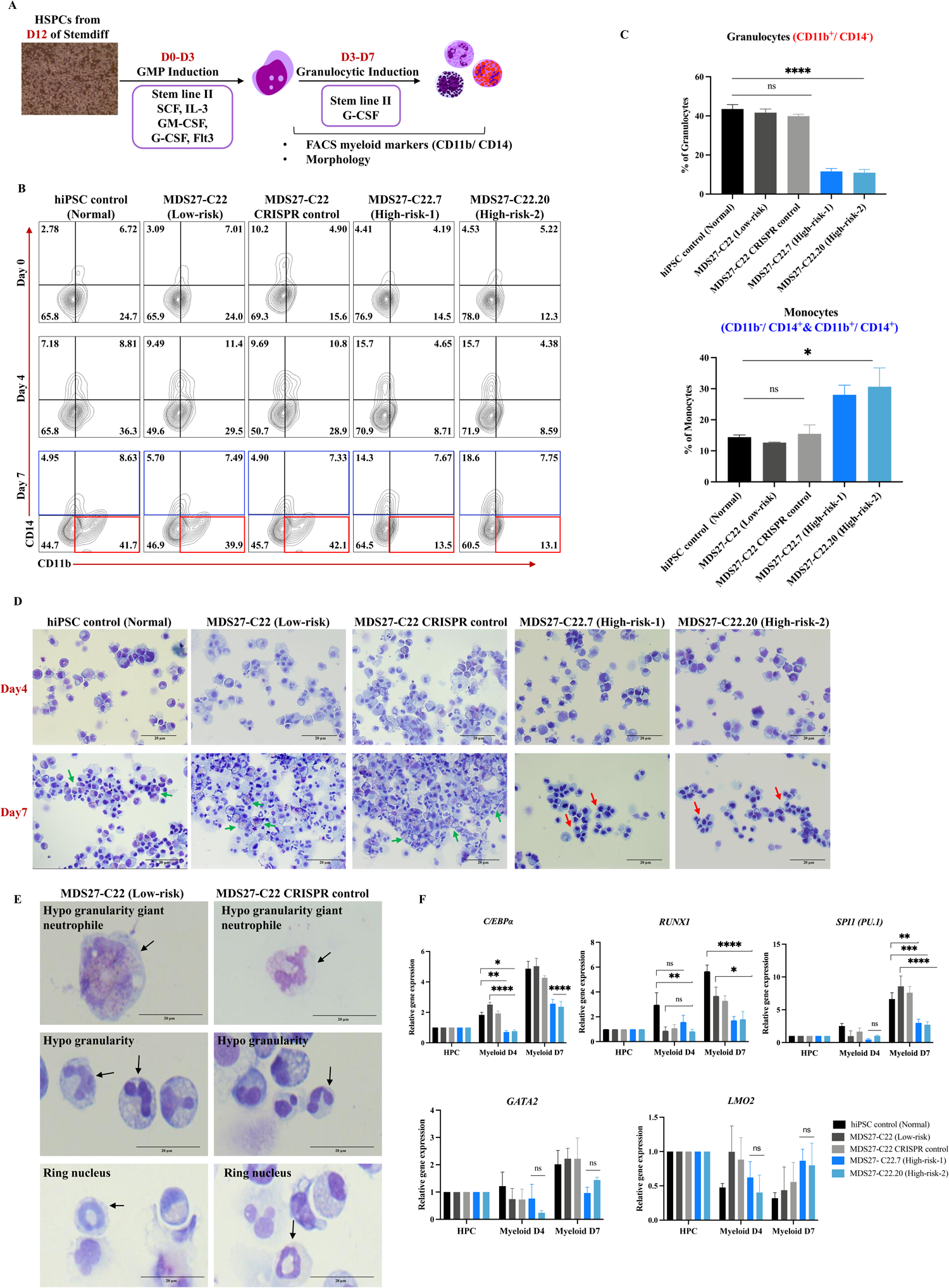
High-risk MDS-iPSC containing a *C/EBPα* mutation exhibit a block in the granulocytic differentiation. (A) Schematic representation the myeloid differentiation protocol. (B) Representative images of flow cytometric analysis of CD11b^+^ CD14^-^ (granulocytes) and CD11b^+^ CD14^+^ (Monocytes) for the indicated cell types on day 0, day 4 and day 7 of differentiation. N= 4 independent experiments. (C) Percentages of CD11b^+^ CD14^-^ (granulocytes) and CD11b ^+^ CD14^+^ (Monocytes). Mean and SEM are shown **** p <0.0001, * p< 0.05, and (ns, no significant), Paired t test. N= 4 independent experiments. (D) Diff-quick stained cytospins showing common the presence of granulocytic cells (green arrows) and erythroblasts (red arrows). The pictures were taken by Leica DM6000 microscope at 40x magnification, 20 μm scale bar. N= 4 independent experiments. (E) Diff-quick stained cytosipin showing aberrant morphology of myeloid cells (black arrows) in MDS27-C22 (Low-risk) and MDS27 C22 CRISPR control. The pictures were taken by Leica DM6000 microscope at 40x magnification, 20 μm scale bar. N= 4 independent experiments. (F) RNA expression of *C/EBPα, RUNX1, SPI1 (PU.1),GATA2 and LMO2* measured by qRT-PCR, normalized against the *GAPDH* housekeeping gene, and expressed relative to HSPCs expression. N= 3 independent experiments.

Morphological examination of the myeloid cells by light microscopy was performed to evaluate distinct maturation features such as granule content, nuclear morphology and cytoplasm/nucleus ratio. Interestingly, the committed eosinophils and neutrophils were identified within the myeloid population of hiPSC control, MDS27-C22 (low risk) and MDS27-C22 CRISPR control hiPSCs, accompanied by appropriate segmented nuclear morphology and granules (Figure 4D, green arrows). However, granulocytes were not detected in the myeloid cultures derived from high risk hiPSCs, in agreement with the flow cytometry data (lack of the CD11b ^+^CD14^-^ population). In addition, morphological assessment of myeloid cells derived from high risk hiPSCs showed that mutant clones differentiated into a heterogenous population that contained mainly erythrocytes (Figure 4D, red arrow). sInterestingly, the morphological results of myeloid cells of *CEBPA* WT lines (MDS27-C22 and MDS27-C22 CRISPR control) revealed dysplastic morphology in granulocytes such as hypo-segmentations, ringed nuclei, and hypo-granularity (Figure 4E). These aberrant morphologies are consistent with those reported previously in MDS patients ^29^.

As expected by the change in the open reading frame, the C/EBPα mutation had significant effects on the functionality of the protein and consequently downregulation of the C/EBPα target genes *CEBPA, SPI1 (PU.1)* was observed in myeloid cells derived from the *C/EBPα ^mut^* iPSC lines F(igure 4F).

Moreover, *LMO2* was shown to be upregulated in all MDS27 iPSC lines (low and high-risk) by day 7 of myeloid differentiation compared to the iPSC control line. As LMO2 has been shown to induce erythroid differentiation ^30^, the increase in *LMO2* expression in our cultures could explain the increase of erythroid cells during the myeloid differentiation (Figure 4F). The qRT-PCR data indicated that the *CEBPA* mutation leads to deregulation of the transcription factor expression signature resulting in the inhibition of myeloid differentiation and induction of transition to the erythroid lineage.

Overall, the *CEBPA* mutation in the context of *ASLX1, RUNX1* and *SRSF2* mutations leads to an inhibition of granulocytic differentiation and a significant increase in the proportion of erythroid lineage cells. However, the original mutations *ASLX1, RUNX1, SRSF2* are responsible for the specific dysplastic morphology observed in myeloid cells.

### Dyserythropoiesis of high-risk MDS-iPSC containing the CEBPA mutation

Next, we wanted to determine if the formation of mature erythrocytes could be affected by the the *CEBPA* mutation. We profiled erythroid differentiation kinetics by measuring the expression of the erythroid markers CD71 and CD235a: from erythroid progenitor cells (CD71 ^+^) to erythroblasts (CD71 ^+^ CD235a^+^) and mature erythrocytes (CD71 ^-^ CD235a^+^). The kinetics of erythroid differentiation in hiPSC control and MDS27-C22 and MDS27-C22-CRISPR control iPSC lines, showing similar percentages of erythrocyte progenitors, erythroblasts and mature erythrocytes at any given time point (Figure 5A).

**Figure 5:**
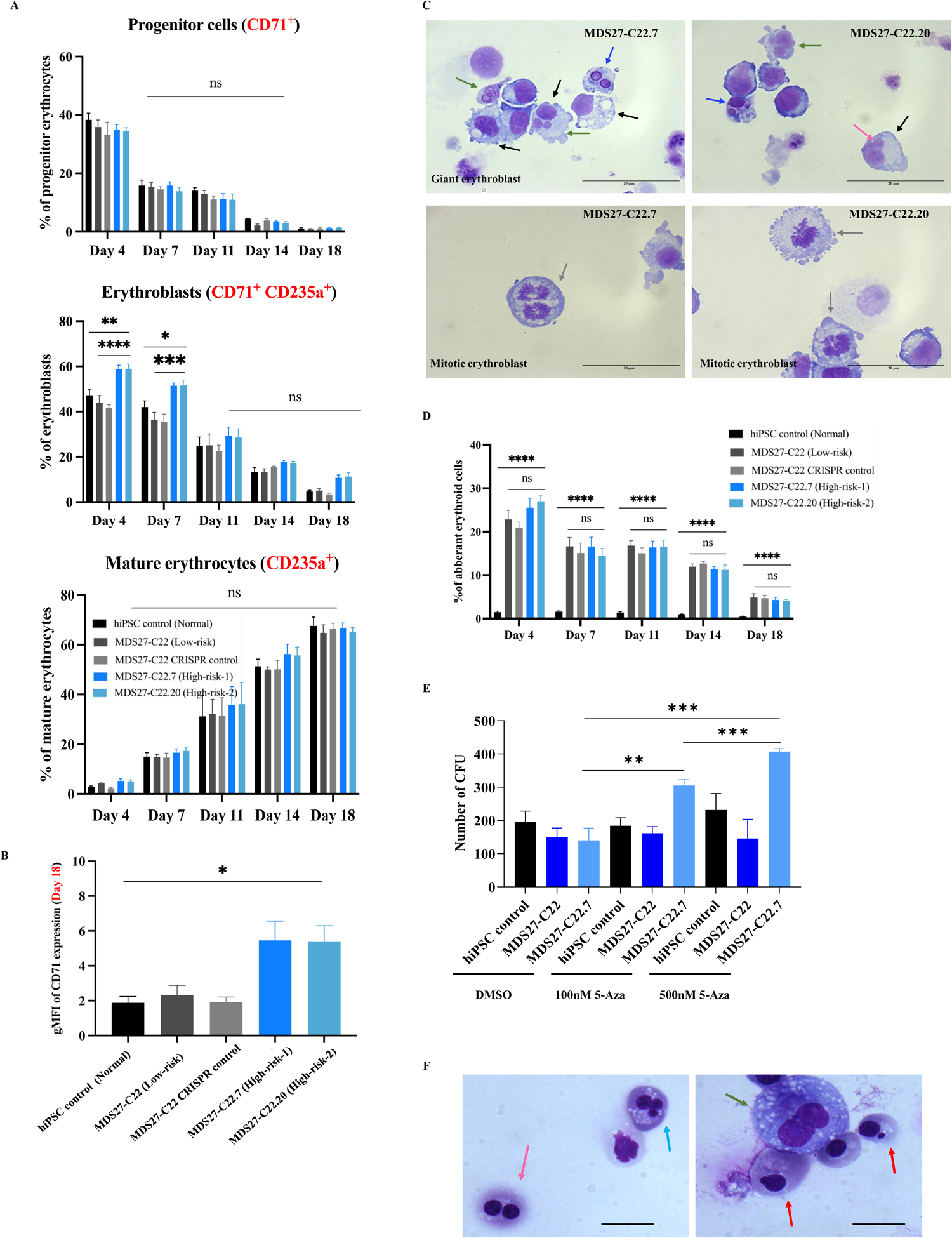
Erythroid differentiation of high-risk MDS-iPSC containing a *C/EBPα* mutation reveals new dysplastic features. (A) Percentage of erythrocyte (CD71 ^+^), erythroblasts (CD71 ^+^ CD235a^+^) and mature erythrocytes (CD235a ^+^) during several time point of erythroid differentiation. Mean and SEM are shown. **** <0.0001, **p<0. 001, * p<0.05 and (ns, no significant), Two-way ANOVA. N= 4 independent experiments. (B) Bar graph showing CD71 geometric mean (gMFI) of each line at day 18 of erythroid differentiation. Mean and SEM are shown. * p<0.05, One-way ANOVA with multiple comparisons. N= 4 independent experiments. (C) Diff-quick stained cytospins showing a new aberrant morphology; Giant erythroblast (black arrow), Mitotic erythroblast (Gray arrows) in MDS27-C22.7 (High-risk-1) and MDS27-C22.20 (High-risk-2), Multinucleated erythroblast (green arrows), bi-nucleated erythroblast (blue arrows) and nuclear bridge (pink arrows). The pictures were taken by Leica DM6000 microscope at 100x magnification, 20 μm scale bar. N= 4 independent experiments. (D) Bar graph represents the percentage of the aberrant cells of MDS27 clones relative to the aberrant morphology of hiPSC control. Results are presented as mean ± SEM and **** p >0.0001, Two-way ANOVA with multiple comparisons. (E) Bar graph represents number of CFUs obtained from control, MDS27-C22 (Low-risk) and MDS27-C22.7 (high-risk) in the presence of different concentration of 5-Aza or vehicle control (DMSO). Mean and SEM are shown. **** p < 0.0001, ** p<0.001, * p 0.05 and (ns, no significant), Two-way ANOVA. N= 3 independent experiments. (F) Diff-quick stained cytospins from colony assay showing aberrant morphology in MDS27-C22.7 (high risk) after 5-Aza treatment; nuclear budding early orthochromatic erythroblasts (red arrow), bi-nucleated pronormoblast (green arrow), multi nucleated basophilic erythroblast (blue arrows) and binucleated orthochromatic (nuclear bridge) (pink arrows). The pictures were taken by Leica DM6000 microscope at 100x magnification, 20 μm scale bar. N= 3 independent experiments.

In contrast, the *C/EBPα^mut^* high riskh iPSCl in esdisplayed agreaterpercent age of erythroblasts (CD71^+^ CD235^+^) at day 4 and 7 of differentiation compared to any of the control cells (Figure 5A) and displayed a higher CD71 expression at the end of the erythroid differentiation (Figure 5B). Interestingly, this result indicates that disruption of the bZIP domain in cells harbouring *ASXL1, RUNX1* and *SRSF2* mutations impaired proper terminal erythroid differentiation as well.

Morphological analysis revealed aberrant morphologies such as giant erythroblasts and mitotic erythroblasts in *CEBPA^mut^* MDS27-iPSC lines but not in MDS27-C22-CRISPR control iPSC, indicating that these new aberrant cells are associated with the acquisition of the *CEBPA* mutation to cells harbouring the *ASXL1, SRSF2* a n d*RUNX1* mutatio (Figures 5C). More over, the percentage of erythroid cells with aberrant morphology of high-risk iPSC compared to control hiPSC was significantly higher during differentiation (Figure 5D). However, the percentage of aberrant cells in the high risk hiPSC lines compared with MDS27-C22 low risk and MDS27-C22-CRISPR control hiPSCs was slightly increased but not statistically significantFi(gure 5D).

Taken together, cells harbouring mutations in the four genes (*SRSF2, ASLX1, RUNX1* and *CEBPA*) display an alter kinetics of terminal erythroid maturation with evident signs of dysplastic erythroid differentiation.

### High-risk MDS-iPSC containing C/EBPα ***^mut^*** are resistant to 5-Azacitidine treatment

Our work thus so far showed that the isogenic iPSC lines generated could replicate the disease phenotype with dysplasia observed in both erythroid and myeloid linages upon HPCs differentiation. Previously, derived iPSCs from MDS patients have been shown to be useful to assess drug response, with high-risk MDS responding to conventional demethylating agents such as 5-Azacytine (5-AzaC) ^31^. The MDS27 patient did not respond to 5-Aza after disease progression, and thus we were interested to determine whether our *in vitro* system could recapitulate therapy resistance. To do so, we concentrations of 5-AzaC (Supplementary Figure 7A). Treatment of 5-AzaC did not have any effect on colony growth in healthy iPSC or low risk iPSC clones (Figure 5E). In contrast, 5-AzaC treatment had a profound effect in cells derived from the high risk MDS, with a massive expansion on colony number, specifically in BFU-E F(igure 5E and Supplementary Figure 7B ad 7C) which was also dose dependent. To determine whether the expansion of the BFU-E colonies was an indication of healthy recovery, we performed a morphological analysis of the cells grown in colony assays. This analysis revealed the presence of dyserythropiesis in the high risk iPCS after 5-Aza treatment (Figure 5F), characterised by binucleated (pink and green arrows, Figure 5F), multinucleate (blue a,rFrioguwre 5F) and budding nuclei (red arrows, Figure 5F) erythroid cells, and thus confirming that cells did not respond to 5-Aza treatment. Taken together, our work shows that the isogenic iPSC lines generated recapitulate the disease phenotype and the therapy resistance observed in the MDS27 patient.

### The CEBPA mutation affects the chromatin landscape of distal cis-regulatory regions

To gain a mechanistic insight into how the *CEBPA* mutation promoted disease progression in MDS, we analyzed open chromatin regions in sorted CD34 ^+^CD45^+^ HSPCs derived from low risk (MDS27-C22) and high risk (MDS27-C22.7) iPSCs, using the assay for transposase-accessible chromatin with high-throughput sequencing (ATAC-seq). Our analysis revealed that the chromatin landscape shifted after the acquisition of *C/EBPα* mutation, with 1691 regions gaining accessibility and 1576 regions losing accessibility (>1.5-fold change) in *CEBPA* mutant HSPCs (open regions in low-risk samples) (Figure 6A). The effect of t*C*h*E*e*BPA* mutation was more pronounced in distal elements (observed 79.3% and 75.9% differential peaks; in MDS-High and Low risk respectively; compared to 59.6% in all peaks; Chi-square p value 5.4E-14) than in promoter regions (Figure 6B). The analysis of motifs specific for open chromatin regions present in low risk HSPCs and lost in high risk HSPC samples showed a strong enrichment of PU.1, CEBP and KLF sites, consistent with the *CEBPA* mutation disrupting the DNA-binding domain and thus preventing the interaction of *CEBPA* with its genomic targets^32, 33^ (Figure 6A and 6C). In contrast, open chromatin regions only present in MDS-high risk HSPCs were enriched in binding motifs for ETS, GATA, AP-1 and RUNX transcription factors (Figure 6A and 6 D); with the presence of the GATA signature, probably indicating a more immature state. Genes with differential accessible chromatin sites included over 100 transcription factor genes, among them *MAF, CEBPE* and *RUNX1*, (Figure 6E). Collectively, our data illustrates that the presence of a single genomic copy of mutant *CEBPA* carrying a mutation in the mid region of the protein alters the chromatin landscape of HPCs, specially of distal cis-regulatory regions.

**Figure 6:**
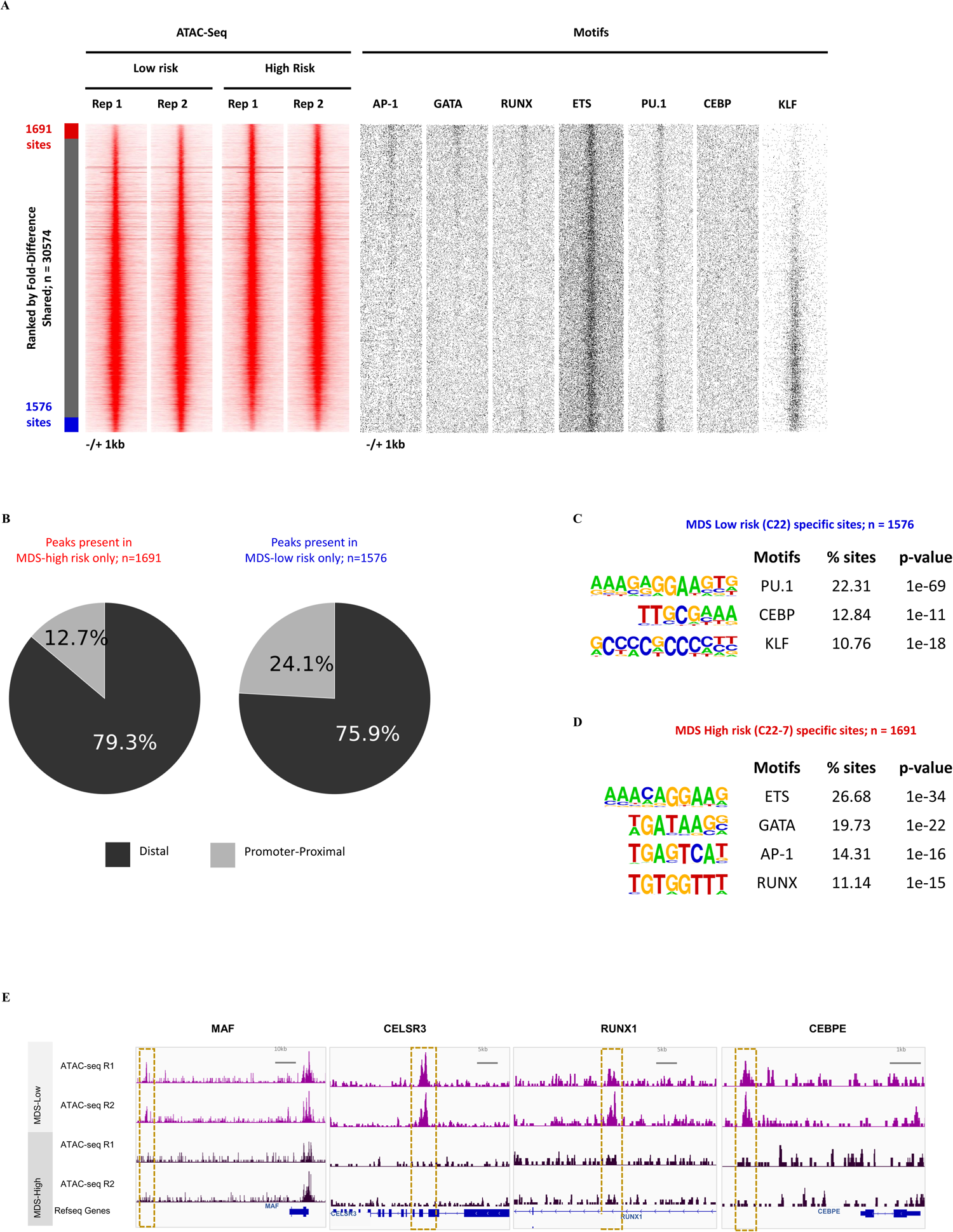
Changes in distal cis-regulatory elements in HSPCs with *C/EBPα mutation*. (A) Profiles of the ATAC-seq signals within each 2000-bp window centered on each peak for Day14 CD34^+^CD45^+^ HSPC-iPSC derived from Low-risk MDS (C22) and from high-risk MDS (C22.7) (N=2 independent experiments). Peaks are shown in order of decreasing log2 fold-difference between Low and High risk samples (see methods). Positions of transcription factor binding motifs are plotted alongside. (B) Pie chart showing percentage of peaks at promoters and distal regulatory elements from peaks present in high-risk MDS HSPCs only (left) or in low-risk MDS HSPCs (right). (C) Motif enrichment analysis (Homer *de novo* motifs) for peaks only present in low risk HSPCs compared to unchanged peaks. (D) Motif enrichment analysis (Homer *de novo* motifs) for peaks only present in high risk HSPCs compared to unchanged peaks. (E) ATAC-seq UCSC genome browser screenshot depicting accessible chromatin sites being differentially regulated between the low- and high-risk HSPCs. Red squares show differential ATAC-seq peak between both conditions. Y-axis are set at 70 RPKM.

### Clonal evolution during disease progression

To determine whether changes in the chromatin landscape associated with the acquisition of the *CEBPA* mutation affected the transcriptional programme, we performed single cell transcriptome analysis in Day 12 HSPCs -iPSCs derived from the BU3.10 iPSC control line, MDS27-C22 (low risk) and MDS27-C22.7 (high risk). We also performed single cell transcriptomics in HSPCs 4 days after differentiation towards the erythroid lineage from BU3.10 iPSC control line and low risk to define transcriptional changes during differentiation (Supplementary Figure 8A).

Unsupervised clustering revealed 13 clusters (Figure 7A and 7B). Clear changes in cluster representation were observed between wild type, low risk and high risk HSPCs (Figure 7A). Also, changes in the cell cycle status was evident between the samples, with lower proliferation rate in the high - risk sample comparetolowrisk, in dicative of amorequies centstage (Figure 7B and Supplementary Figure 8B). Based on gene expression markers, wild type HPCs could be found mainly in three clusters: 2, 9 and 4 which were identified as early stem/progenitors (HPCI) (*HMG high, GYPB*), early erythroid/megakaryocyte progenitors (MEP) (GP9, GP1BB, Rap1b, PF4) and myeloid progenitor cells (GMP/monocytic) (*MPO, AZU1*), respectively (Figure 7C, Supplementary Figure 8C-D)^34^. A prominent early progenitor cluster was also present in the MDS low risk sample (HPCs II, cluster 1), expressing similar markers as the wild type HPC cluster (*MYB, NPM1, GATA2, HMGA* and *GYPB),* but this cluster was really underrepresented in the high risk MDS comprising of 6% of the total cell population as compared to 60% in low-risk MDS population (Figure 7C-D and Supplementary Figure 8D). MDS clusters 3, 6, 7, 8, 11 and 12 were distinct from wild type. A gradual loss of cell identity from low risk to high risk could be observed representing a unique MDS/AML cell populations, with 30% of the cells in low-risk forming part of the unique MDS/AML signature compared to 90% of the cells in high-risk. Additionally, changes in the size of these clusters were noticeable, indicating clonal dynamics during disease progression. Hence, some of the clusters were greatly expanded from low to high risk MDS (red squares, Figure 7D), whereas others were reduced from low to high risk MDS or did not show a clear difference in composition (blue squares and green square, respectively, Figure 7D).

**Figure 7:**
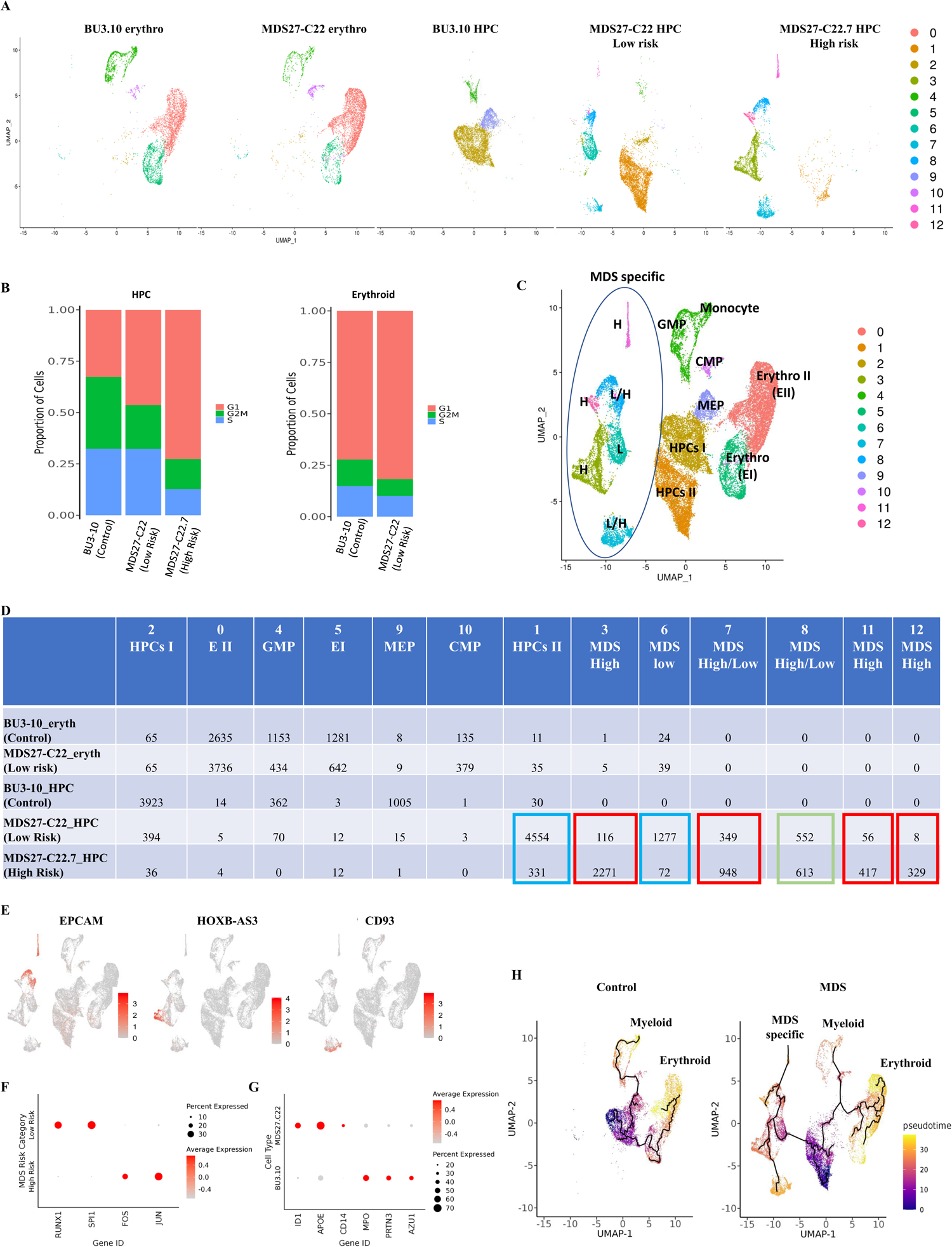
Distinct MDS transcriptome signature and clonal evolution during disease progression measured by single-cell transcriptome analysis. (A) Analysis of scRNA-Seq data. Uniform Manifold Approximation and Projection for Dimensional Reduction (UMAP) map for individual samples. Each dot in the map represents a cell and is colored accordingly to cluster assignment. (B) Histogram showing the proportion of cells in each cell cycle phase within each cluster as identified by the expression of cell cycle regulated genes. (C) 13 clusters based on expression of specific genes. Cluster annotation was based on manual curation of marker genes. Each dot in the map represents a cell and is colored accordingly to cluster assignment. (D) Table representing the number of cells in each cluster. Blue squares, clusters reduced during disease progression; red squares, clusters increased during disease progression; green squares, no change during disease progression. (E) Expression of indicated genes projected on the UMAP map. Colour intensity represents expression data log2 normalized unique molecular identifier (UMI) counts. (F) Dot plots showing the scaled expression level of the indicted transcription factors in low risk and high risk MDS samples. Colours represent the scaled expression and size encodes the proportion of gene-expressing cells. (G) Dot plots showing the scaled expression level of the indicted myeloid genes in control and low risk samples. Colours represent the scaled expression and size encodes the proportion of gene-expressing cells. (H) Monocle pseudo-time trajectory for MDS specific, myeloid and erythroid cells projected on the UMAP map of scRNA clusters. Cells are coloured according to their pseudo-time value.

Among the most highly expressed genes in the MDS clusters were genes associated with AML as for example, *CD99* ^35, 36^ *, EpCAM,* related to enhanced cell survival, diminished mitochondrial metabolism, ribosome biogenesis, and differentiation capacity and an activated transcriptomic signature associated with AML ^37^ (Figure 7E and Supplementary Figure 8D), *ID1,* which is expressed by AML cells and has been shown to increase leukemic proliferation and promote tumour progression by impairing myeloid differentiation ^38, 39^, *ID3*, which promotes erythroid differentiation^40, 41^ (Supplementary Figure 8D). Other leukemia associated genes were expressed by cells within the clusters expanded during disease progression, for example *HOXB-AS3* (Figure 7E), a gene encoding a long non-coding RNA shown to be an adverse prognostic marker for AML and MDS^42^, *CD93* (Figure 7E), a marker associated with proliferative leukemic stem cells ^43^ and genes associated with angiogenesis and therapy resistance (*RAMP2, CALCRL*)^44, 45^, angiogenesis and poor prognosis in MDS (*FLT1, KDR*)^46, 47^ (Supplementary Figure 8D).

To gain a better insight in the changes occurring during disease progression, we combined all the cells from low risk and high risk into a “pseudo-bulk” sample and performed a differential gene expression analysis. KEGG pathways analysis of the approximately 500 differentially expressed genes (Supplementary Table 2) indicated that low-specific genes were related to a myeloid-differentiation type signature whilst genes expressed in high risk MDS cells encoded for cell signalling components, such as Wnt signalling and Hippo pathway, both associated with AML (Supplementary Figure 9A-B). Additionally, a clear difference in the expression of important transcription factors regulating hematopoiesis was noticeable between low risk and high risk HSPCs samples, with high-risk cells showing reduced expression of *RUNX1* and *SPI1* (*PU.1*) and gaining expression of the AP1 factors *FOS* and *JUN* (Figure 7F, Supplementary Figure 8D).

Once HSPC were subject to erythroid differentiation, cells from control and MDS low risk were both found in clusters which represented different stages of erythroid (clusters 0 and 5) and myeloid (clusters 4 and 10) differentiation. A different representation of myeloid and erythroid cells between control and MDS low risk samples was evident, with less myeloid and more erythroid cells in MDS low risk compared to control (Figure 7A and 7D). By inspecting the most highly expressed genes within each cluster, we found clear differences in gene expression between the control and MDS low risk samples. For example, *ID1, APOE* and *CD14* were expressed by MDS low risk cells from the myeloid cluster whereas *MPO, PRTN3* and *AZU1* genes were expressed by wild type BU3.10 cells, revealing changes in myeloid composition between control and MDS low risk cells (Figure 7G, Supplementary Figure 8D and Supplementary Table 3).

To gain more insight into the position of the aberrant cell population within the differentiation trajectory, we performed a pseudo-time analysis (Figure 7H). The control cells showed a clear trajectory, with myeloid progenitors (left) and erythroid progenitors (right) branching off from the more immature HSPCs (purple/blue). In clear contrast, MDS cells showed an additional trajectory branching off from the immature HSPCs, with branch points corresponding to the myeloid lineage and a disorganised erythroid lineage (Figure 7H). These results are consistent with our differentiation results, and an increase in erythroid differentiation at the expense of the myeloid differentiation.

In summary, the single cell transcriptome analysis is consistent with the notion of MDS disease arising from an early stem/progenitor cell, leading to dysregulation of genes involved in stem/progenitor cell development which is translated into a different clonal composition, loss of cell identity and expansion of specific clones during disease progression. Our isogenic model system of disease progression shows changes in clonal composition and emergence of new clones which appear during disease progression as a consequence of the acquisition of a C/EBPα mutation.

## DISCUSSION

The extensive heterogeneity of MDS is the result of a combination of different mutations that blood progenitor cells acquire during ageing, leading to clonal evolution and disease progression. In patients, mutations in components of the spliceosome, and epigenetic modifiers can act as driver mutations as well and when occurring together generate a combination that is causal for MDS development^5^. Through PCR and sequencing-based studies, mutations in *C E B P A*have also been shown to associate with disease progression and AML ^12, 48, 49^. However, direct proof for this idea has so far been lacking as it requires to systematically test the contribution of different mutations to MDS pathogenesis in an isogenic background.

In the work described here, we applied somatic reprogramming to iPSC and CRISPR technology to generate human isogenic MDS cell lines from patients before and after disease progression. These experiments identify the disruption of CEBPA bZIP domain as causative for disease progression in MDS, affecting cell fate decision in the evolution from low risk to high risk. Moreover, by disruption of the CEBPA bZIP domain in wt and mutant background, we were able to dissect the contribution of different mutations to the disease phenotype. We show (i) that the heterozygous disruption of the bZIP domain in a wt background leads to an impediment of granulocyte differentiation, but does not confer self-renewal capacity (ii) that the primary mutational pattern (*ASXL1, SRFS2* and *RUNX1*) leads to dysplasia of erythroid and myeloid lineages, and (iii) that the additional introduction of the mutation causing the disruption of the bZIP domain leads to a combination of both, and promotes self-renewal capacity of HPCs, thus mimicking the increased risk of AML development seen in the patient.

Our findings are broadly consistent with the pathology data reported for most MDS patients, including a low number of CFUs progenitor cells ^8, 31, 50–52^. Being able to follow aberrant blood cell development of cells with a different mutational background in liquid culture revealed the development of mature cells with aberrant morphology, such as nuclear bridges and multinuclear content which are common dysplastic features of MDS patients ^29, 53^. Moreover, we were also able to recapitulate the therapy resistance to 5-AzaC as seen in the patient. Our MDS27-iPSCs model is therefore a *bonafide in vitro* model for early MDS disease pathology showing erythroid and myeloid dysplasia similar to the anaemia and cytopenia of the patient at the time of diagnosis and after disease progression.

Our work also shines an interesting light on the role of mutant C/EBαPand its interaction with other mutations in MDS pathology when all mutations are present as heterozygotes. Similar percentages of HPCs were produced by the isogenic low- and high-risk clones. In agreement with the essential role of C/EBPα in the formation of granulocyte progenitors ^54, 55^, neither CFU-G nor mature granulocytes were detected in colony assays or liquid cultures, respectively. Nonetheless, the high-risk iPSC had the potential of forming CFU in methylcellulose, in much higher numbers than those reported previously ^8^. It has been reported that *CEBPA* deleted fetal liver progenitors are hyperproliferative, fail myeloid differentiation, and show increased self-renewal potential *in vitro and in vivo* ^55, 56^. This data precedes published results where mouse bone marrow mononuclear cells transduced with a C-terminal mutated C/EBPα (*CEBPA^Cmut^*) increased the self-renewal of the CFUs forming colonies after 6 rounds of re-plating ^48^. Moreover, 5-FU treated mice transduced with these *CEBPA^Cmut^*bone marrow cells developed AML when transplanted into recipient mic^4^e^8^. However, our data suggest that in a wild type background, a frameshift mutation in the mid region of *CEBPA* did not confer self-renewal capacity. Thus, the increase in self-renewal capacity after the acquisition of *CEBPA* mutation must be due to a synergistic effect with other mutations present within the clone. It has been previously reported that RUNX1 mutations enhance self-renewal and impede granulocytic differentiation^57^. Thus, it is reasonable to speculate that the addition of the *CEBPA* mutation affecting the bZIP domain further imposes a granulocytic differentiation impediment and together with RUNX1 mutation promotes an increased proportion of erythroid progenitors at the expense of myeloid progenitors.

In spite of the presence of a wild-type allele, our data show that the presence of a C/EBP α protein with a frameshift mutation in the mid region of the protein affects the chromatin landscape through dysregulation of accessible chromatin regions in distal cis-regulatory elements, suggesting a dominant deregulatory activity of this protein, probably by interfering with dimerisation. Lost chromatin sites were enriched in PU.1, KLF and CEBPA motifs, suggesting that C/EBP α binding was lost which agrees with the change in open reading frame of the mutated protein. High risk HPCs gained ATAC-seq peaks enriched in ETS and RUNX motifs, which have been reported to regulate the leukemic signature in AML patients with CEBPA biallelic mutations (*CEBPA^N/C^*)^58^, whilst enrichment of GATA motifs indicates that the HPCs from high risk are less differentiated than those from low risk, which is consistent with the absence of myeloid differentiation.

By single-cell mutational studies, it has been reported that changes in clonal architecture occur during MDS disease progression ^59, 60^ and single-cell transcriptomics for paired-samples from two patients that evolve from MDS to AML revealed changes in genes associated to signalling pathways such as TGF-β and TNF α^61–63^. Our isogenic iPSC lines derived harbouring the genetic make-up of MDS27 patient allow us for the first time to study clonal evolution and the transcriptional changes imposed by heterozygous disruption of the C/EBP α bZIP domain during disease progression as measured by single cell RNAseq. Changes in the size of MDS specific clusters was observed between low and high-risk samples and important genes previously associated with AML were expressed by cells within these clusters such as *ID1, CD93* and *CALCRL*. Genes coding for several insulin-growth factor binding proteins (IGFBP) were expressed by cells in different MDS clusters. For example, *IGFBP3*, associated with early stage of MDS ^64^, was expressed by cells in low risk but not by cells in high-risk clusters. Additionally, our data showed an upregulation of genes encoding for important signalling pathways associated with MDS progression and AML such as Wnt and TGF β signaling^65–68, 62^. Our single cell RNAseq analysis also revealed that disruption of C/EBP α bZIP domain affects the cell cycle of the cells, with a block in G1 as expected for cells with a differentiation block.

C-terminal mutations on the bZIP domain (*CEBP*A*^bZIP^*), are frequently in frame mutations associated with a favourable prognosis *d*in*e novo* AML ^15, 16^ and recently have been classified as a distinct AML category by WHO. Our work suggests that this classification could be expanded to frameshift mutations within the mid region that would equally disrupt the bZIP domain. Importantly, our isogenic iPSC lines provide a bona fide *in vitro* experimental system that could be utilised as a platform for drug screening and for future studies regarding the molecular mechanisms of drug resistance.

## Acknowledgments

The authors wish to thank Dr Giacomo Volpe for critical reading of the manuscript. We also thank Genomics Birmingham for NGS sequencing and Mary Clarke for cell sorting. RA and YA PhD studentships were funded by the Ministry of Education and Royal Embassy of Saudi Arabia Cultural Bureau. This work was also funded by a FIS grant ISCIII (PI19/00730) to EB.

## Author Contributions

Methodology, R.A., Y.A., R.B.; Formal Analysis, P.K., C.W., I.A., T.S.; Investigation, R.A., Y.A., C.S., E.B., T.S., A.F., S.K., M.B.; Resources, G.J.M., P.M., M.R.; Writing-Original Draft, P.G.; Writing- Review & Editing, P.G, C.B., S.K., E.B., G.J.M., P.M; Funding acquisition, P.G. and E.B; Conceptualization, P.G; Supervision, P.G.,C.B; Project Administration, P.G.

## STAR Methods

### iPSC generation

#### Human iPSCs reprogramming using the integration-Free Sendai virus

Human iPSCs from MDS27 were generated from the peripheral blood sample collected in 2013. MDS27 cryopreserved peripheral blood mononuclear cells were thawed and cultured in expansion medium in a 24-well plate for 3-6 days. At day 6, the cells were counted and 25×10 ^4^ cells were resuspended in 300 fresh expansion media with 4μg/ml Polybrene (Sigma, TR-1003) and placed in a 15ml falcon tube for transduction, MOI 3 of Sendai Virus containing the four Yamanaka factors was added to the cells for three hours at 37 °C. Two days later, the transduced cells were centrifuged at 300g for 8 minutes and plated in a 6-well plate containing 90%-confluent of mitomycin C-treated MEFs and “day 8 medium” for three days at 37 °C, and every two days the medium was changed. At day 11, the medium was replaced with “day 11 medium” and incubated for four days at 37 °C and every two days the medium was changed with “day 11 medium”. At day 16, the medium was changed to hiPSC medium and every day thereafter, the medium was changed to fresh “hiPSC medium”. At day 25, emerging iPSC colonies were picked individually into a 6-well plate coated with Matrigel hESC-qualified Matrix (corning life science, 354277). The colonies were cultured in hiPSC medium with 10 μM of Rock inhibitor (Y-27632, LKT laboratory, Y1000) and incubated at 37 °C. From the first day after picking onwards, the colonies were cultured continuously using hiPSC medium without Rock inhibitor.

#### Human iPSCs generation by integration-Free episomal reprogramming

The episomal reprogramming of MDS27 PBMNCs from the sample obtained in 2013 was performed as previously described ^9^.

#### AP staining

AP staining was performed according to standard protocols, as previously described ^69^.

#### Immunofluorescence for cytoplasmic marker TRA1-81

Cells were washed twice with StemFlex medium and incubated with TRA1-81 mouse primary Ab for one hour at 37 °C. After three washes with StemFlex medium, the secondary Ab goat anti-mouse-IgM-Alexa, Fluor 488 was added to the cells and incubated for one hour at 37 °C. Then, cells were washed with PBS and fixed with 2% (v/v) Formaldehyde methanol free (PFA, Thermo Scientific, 28906) for 10 minutes at room temperature. Cells were washed twice, mounted and imaged under a fluorescence microscope (Leica DM6000, Leica Microsystems).

#### Immunofluorescence for the nuclear markers NANOG and SOX2

Cells were fixed with 4% PFA (v/v) in PBS for 20 min at room temperature. Then, the cells with aldehyde groups quenched for 10 minutes with 50mM NH 4Cl (Sigma, 254134) in PBS at room temperature. Afterwards, the cells were permeabilized with 0.5% (v/v) Triton X-100 (Sigma, 11332481001) in PBS for 15 minutes at room temperature and blocked with PBS containing 1% (w/v) bovine serum albumin (BSA, Sigma, A1933) + 0.3% (v/v) Triton X-100 + 10% (v/v) FBS+ 1% (v/v) goat serum (Sigma, G9023) for 1hour at room temperature. Then, the cells were incubated with diluted primary mouse antibodies SOX2 and NANOG in blocking buffer for 1hour at room temperature. After the primary antibody, the cells were washed for 30 minutes with PBS + 0.1% (v/v) Tween-20 (Sigma, P9416), and goat anti-mouse secondary antibody was applied to the cells for 1hour at room temperature. Finally, the cells were washed with PBS, mounted, observed, and imaged under a fluorescence microscope (Leica DM6000, Leica Microsystems).

#### Trilineage differentiation of human iPSC

Differentiation towards the three germ layers was assessed using the STEMdiff Trilineage differentiation kit (Stem cell technology, 05230) performed as described by the manufacturer.

#### Hematopoietic differentiation using Stemdiff protocol

Stemdiff protocol (Stem cell technology, 05310) was followed to differentiate hiPSCs to hematopoietic progenitor cells as described by the manufacturer.

#### Clonogenic progenitor assay

10,000 Hematopoietic cells from day 12 of stemdiff differentiation were plated in 35-mm plastic dishes using 1.2 ml per dish of complete MethoCult H4435 medium in duplicate. The plates were incubated at 37 °C, 5% CO2 for 14 days. Colonies were scored after 14 days of incubation based on the morphological criteria as erythroid colonies (CFU-E), granulocyte, erythrocyte, macrophage, megakaryocyte colonies (CFU-GEMM), granulocyte/macrophage colonies (CFU-GM), granulocyte colonies (CFU-G), and macrophage colonies (CFU-M).

#### Erythroid differentiation

1-2 ×10 ^5^/ml Day 10 hematopoietic Stemdiff cultures were plated in a 6-well plate and cultured for 18 or 20 days using the media with different cytokines according to published protocol ^70^. The erythroid differentiation was monitored every 4 days by flow cytometry analysis of erythroid markers (CD71-APC, glycophorin A (CD235a-PE)) and DRAQ5-APCCy7 (staining for nucleated erythrocytes, ThermoFisher, 65088092). Morphological analysis of erythrocytes was assessed after Kwik-Diff staining (Thermofisher, 9990700).

#### Myeloid differentiation

1 ×10^5^/ml Day 12 hematopoietic Stemdiff cultures were plated in a 12-well plate and cultured for 7 days in myeloid differentiation media ^71^. The myeloid differentiation was monitored on day 4 and day 7 by flow cytometry analysis of myeloid markers (CD14-APCCy7 and CD11b-PECy7) (Table 2). Morphological analysis of myeloid cells was assessed after Kwik-Diff staining.

#### CRISPR-Cas9 to introduce C/EBPα mutation into MDS-hiPSC (C22)

A sgRNA targeting C/EBPα gene on chromosome 19 at position 13.11 was designed using an online tool (Trust Sanger Institute Editing database). sgRNA was first phosphorylated, annealed and cloned into pSpCas9 (BB)-2A-GFP (PX458) (Addgene, 48138) following the protocol provided by ^72^. P3 amaxa kit (Lonza, V4XP-3024) was used to necleofect iPSC (programme DS-150). GFP ^+^ cells were sorted 24 hours after using BD FACSAriaTM Fusion (BD bioscience). Seven days post sorting, 24 clones were picked and expanded to extract genomic DNA for assessing the gene editing efficiency. To evaluate nucleofection efficiency, the target region of C/EBPα was amplified using Q5 High-Fidelity DNA polymerase (NEB, M0491) and two designed primers used for PCR: C/EBPα forward 5’GGCCTCTTCCCTTACCAGCC’3 and C/EBPα reverse 5’CTGGTCAGCTCCAGCACCTT’3. PCR products were processed using a T7 endonuclease I (T7EI) assay using the Alt-R genome editing detection Kit (IDT, 1075932) according to the manufacturer’s protocol.

#### qRT-PCR

RNA extraction was performed with Trizol reagent (15596026, Invitrogen) from HSPCs from day12 of stemdiff differentiation and myeloid cells from day4 and day7 of myeloid differentiation. qPCR was carried out in a Stratagene Mx3005P (Agilent Technologies) using Taqman primers. Ct values were calculated and generated by MxPro 3000 Stratagene software. Different gene relative expression values were calculated against *GAPDH* using the ΔΔCt mathematical model ^73^.

#### ATAC-seq

Fragment transposition ATAC libraries were generated following the Omni-ATAC protocol from ^74^. 50×103 viable cells were pelleted in a fixed angle centrifuge at 500xg at 4°C for 5 min, resuspended in 50 μL of cold ATAC-Resuspension Buffer (RSB) (1M Tris-HCl pH 7.4, 5M NaCl, 1M MgCl2 in sterile H2O) containing 0.1% NP40 (Sigma-Roche), 0.1% Tween-20 (Sigma-Roche), 0.01% Digitonin (Promega) and incubated in ice for 3 min. Lysis was washed out by adding 1 mL of cold ATAC-RSB containing 0.1% Tween-20 (Sigma/Roche) only. Nuclei were pelleted at 500xg at 4°C for 10 min and resuspended in 50 μ Loftrans position mix (25 μ Lof 2 x TD buffer (Illumina), 2.5 μ Lof Tn 5 Transposase enzyme (Illumina), 16.5 μL of PBS, 0.5 μL of 1% Digitonin (Promega), 0.5 μL of 10% Tween-20 (Sigma-Roche), 5 μL of H2O). The mixture was then incubated at 37°C for 30 minutes in a heated block shaking at 1000 RPM. This reaction allowed the Tn5 transposase to simultaneously fragment and tag the chromatin with sequencing adapters. Transposed fragments were purified by using the MinElute Reaction Cleanup Kit (QIAGEN) following manufacturer’s instructions and eluted in 21 μL of H2O. The whole eluted product was amplified for 5 cycles (Table 2.5 and 2.6). Pre-amplified library was stored in ice. Reagent Volume (μL) 29 Primer Ad1 25 μM 2.5 Primer Ad2 25 μM 2.5 NEBNext Master Mix 2x (NEB) 25 Transposed Sample 20 PCR reagents for the pre-amplification of transposed fragments Temperature (°C) Duration Cycles 72 5 min 98 30 sec 1 98 10 sec 63 30 sec 5 72 1 min 4 ∞ PCR conditions for the pre-amplification of transposed fragments. To determine the number of additional cycles needed for an optimal library amplification, 5 μL of pre-amplified library were used to set up a RT-qPCR reaction (Table 2.7 and 2.8). Linear relative fluoresce values were plotted against cycles, and the number of additional cycles needed was defined as the cycle number corresponding to 1/3 of the maximum fluorescence intensity (7). Reagent Volume (μL) Sterile H2O 3.76 Primer Ad1 25 μM 0.5 Primer Ad2 25 μM 0.5 SYBR Green 25x _(in DMSO) 0.24 NEBNext Master Mix 2x (NEB) 25 Transposed Sample 20 30 Setup of the RT-qPCR to determine the number of additional cycles of ATAC library amplification Temperature (°C) Duration Cycles 98 30 sec 1 98 10 sec 63 30 sec 20 72 1 min 4 ∞ RT-qPCR conditions to determine the number of additional cycles of ATAC library amplification The remainder of pre-amplified library was amplified for the additional cycles required following the condition in table 2.8. The reaction was purified using the QIAquick PCR Purification Kit (QIAGEN) following manufacturer’s instructions. Briefly, the sample was mixed with 5 volumes of Buffer PB and loaded onto a MinElute column (QUIAGEN). The column was centrifuged and washed with 750 μL of Buffer PE. Two sequential centrifugations were performed to remove any residual ethanol. All centrifugation steps were performed at 17900 x g for 1 min. Libraries were eluted in 20 μL of H2O and, to avoid adapter contamination, further purified by adding 1.2x volumes of AMPure XP beads (Beckman Coulter) following manufacturer’s instructions. The libraries were then eluted in 20 μL of H2O. Sequencing was performed using a NextSeq2000 to obtain a minimum of 50 million reads per sample.

Raw sequencing reads were processed to remove low-quality bases and Nextera ATAC adapter sequences using Trimmomatic v0.39 ^75^. Reads were then aligned to the human genome (version hg38) using Bowtie2 v2.2.5 ^76^ with the --very-sensitive-local parameter. Potential PCR duplicates were removed from the alignments using Picard MarkDuplicates v2.16.10 (http://broadinstitute.github.io/picard). Peaks were called using MACS2 v2.2.7.1 ^77^ with the options --nomodel --call-summits -q 0.05 -B --trackline. The resulting peaks were then filtered to remove peaks found in the hg38 blacklist ^78^ and to retain only peaks that had a summit height greater than 5 in both replicates of either low- or high-risk MDS clones. Peaks were then combined to produce a single peak union using the merge command in BedTools v2.30.0 ^79^. Read counts were obtained using featureCounts v2.0.1 ^80^ and normalized using edgeR v3.36.0 ^81^ in R v4.1.2. Differentially accessible peaks were identified using the voom method in the Limma package v3.50.3 ^82^. A peak was considered to be differentially accessible if it had a fold-difference of at least 1.5 between conditions. Motif enrichment analysis was carried out in the sets of differentially accessible peaks using the findMotifsGenome.pl function in Homer v4.9.1 ^83^ with the options -size 200 -noknown. To create read density plots, peaks were first ranked according to their fold-difference between low- and high-risk MDS cells. Read densities were then calculated in a 2kb window centered on the peak summit using the annotatePeaks.pl function in Homer with the options -size 2000 -hist 10 -ghist - bedGraph and the bedGraph files produced by MACS2. These were then plotted as a heatmap using Java TreeView v1.1.6r4 ^84^.

#### Single cell RNA sequencing

Hematopoietic cells from the BU3-10 hiPSC control, MDS27-C22 (low risk) and MDS27-C22.7 lines were collected on day 12 of hematopoietic differentiation culture. Erythroid cells were collected on the day 5 of the erythroid culture from BU3-10 hiPSC control and MDS27.C22. Freshly collected cells were passage through columns for cell removal to achieve a viability of >90%. Cells were resuspended in PBS/0/04% BSA (PBS without EDTA and Mg). 100 µl of single cell suspension at concentration of 700-1200 cells/µl (1.2×10^5^/100µL) was submitted to the sequencing service. Library was performed according to the manufacturers instruction (3’ scRNAseq v3.1 10x Chromium) and samples were sequencing using Illumina NextSeq 500. Samples were sequenced at an average of 50,000 reads per cell.

Reads from single-cell RNA-Seq experiments were aligned to the human genome (version hg38) and quantified using the count function in CellRanger v4.0.0 from 10x Genomics and using gene models from Ensembl as the reference transcriptome. Seurat v4.1.0 in R v4.1.2 was used to merge individual data and perform quality control. Cells with greater than 20 percent UMIs aligned mitochondrial transcripts and cells with less than 500 or more than 5000 genes were remo normalized and scaled and merged to form a single dataset. Plotting was done using Seurat functions.

Single-cell trajectory analysis was carried out using Monocle3 v0.2.3 ^85^. Processed data from Seurat was imported into Monocle using the as.cell_data_set command in SeuratWrappers (https://github.com/satijalab/seurat-wrappers). Trajectories were then inferred using the learn_graph command in Monocle. Pseudotime was calculated using the order_cells command, using the earliest inferred HSC population as the root node. These were then plotted using the UMAP coordinates calculated by Seurat.

## Statistical analysis

The statistical analysis was performed using GraphPad Prism 8 (GraphPad Prism version 8.0 for Mac, GraphPad Software, San Diego California USA). All data are expressed as Mean ± Standard Error of the Mean (SEM). *P* values less than 0.05 were considered statistically significant and a star (*) was labelled in the figures. The following statistical analyses were used: Two-way ANOVA to analyse data describing two factors across multiple parametric groups. One - way ANOVA to an describing one factor across multiple parametric.

## Data and software availability

The accession number for the ATAC-seq and single cell RNAseq reported in this paper is GEO:GSE236710.

## Supplementary Figures/Tables

**Supplementary figure 1:**
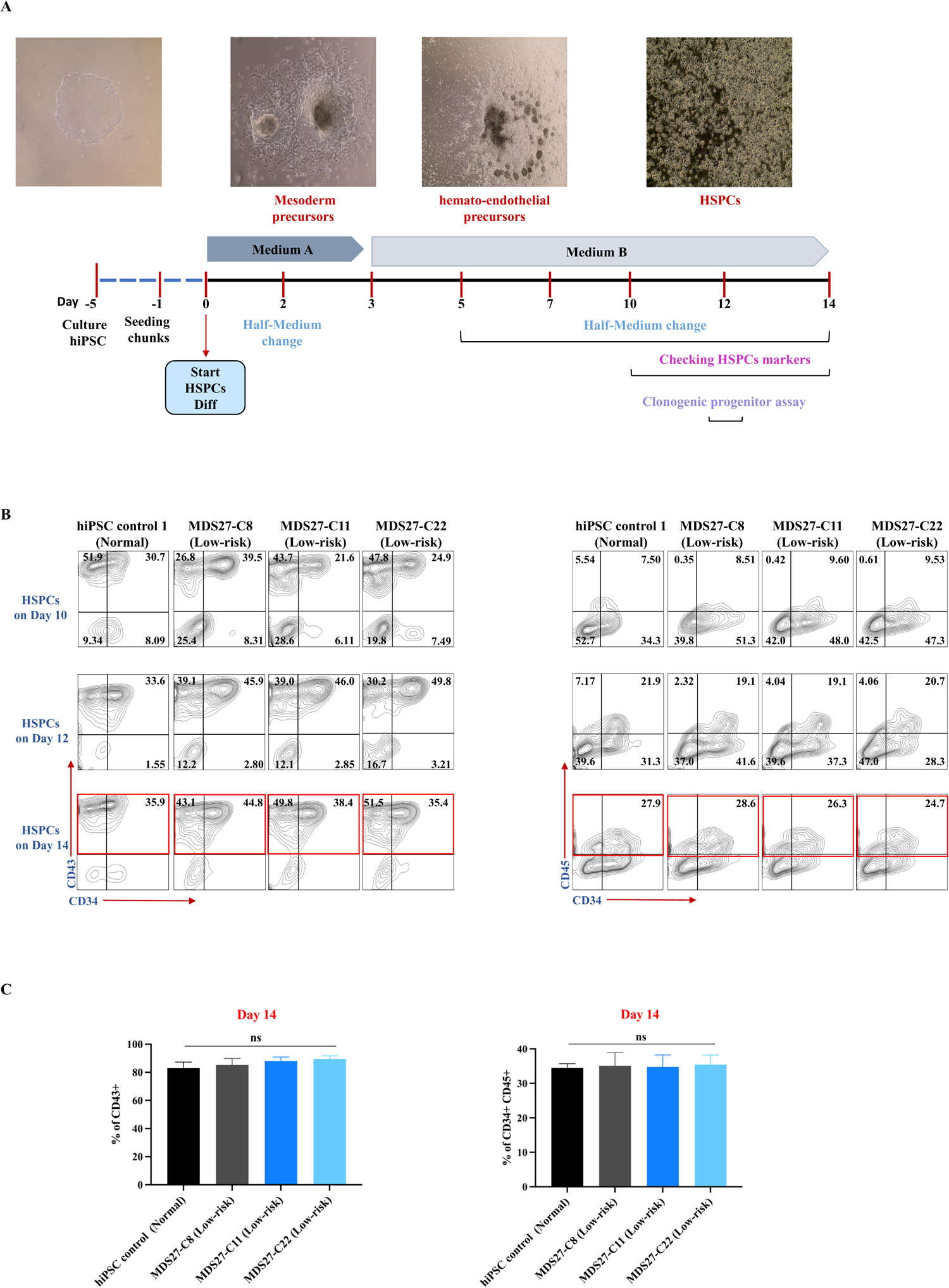
Mutations in MDS27-hiPSCs clones do not affect the HSPCs differentiation. (A) Schematic representation of the HSPC differentiation using hematopoietic STEMdiff protocol from stem cell technology. (B) Representative images of flow cytometric analysis of CD43^+^ and CD34 ^+^ CD4 5^+^ at different times during heamatopoietic differentiation (Day 10, 12 and 14). N=4 independent experiments. (C) Bar graph showing the percentage of early hematopoietic population (CD43 ^+^) and late hematopoietic population (CD34 ^+^ C D 4^+^)5on day 14 of differentiation. Statistical results are presented as mean ± SEM. Ns= no significant, One-way ANOVA with multiple comparisons. N=4 independent experiments.

**Supplementary figure 2:**
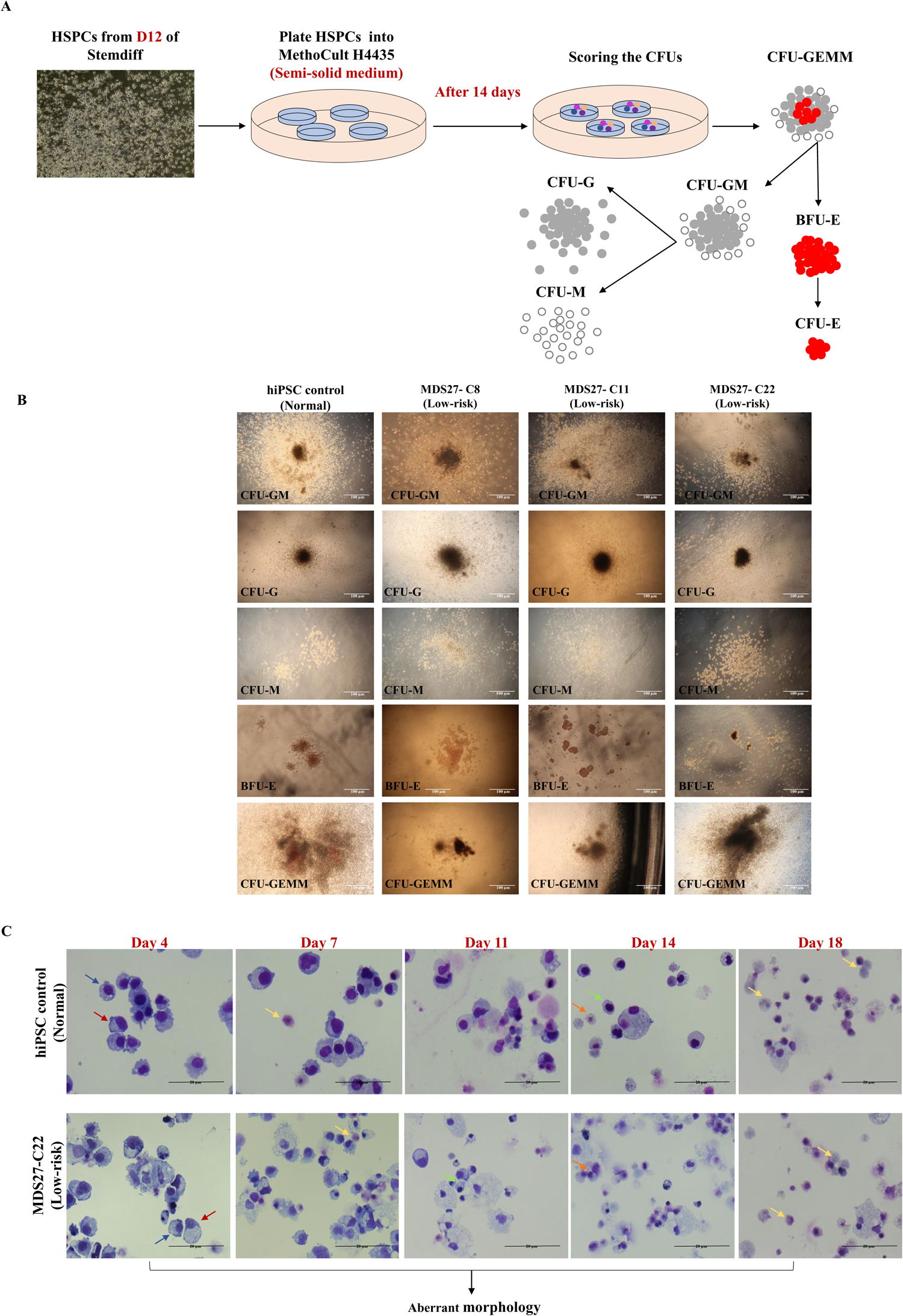
Differentiation potential of HSPCs. (A) Schematic representation of the experimental procedure for the colony forming assays. (B) Representative images of the morphology of the different CFU scored at 14 days in semi-solid medium. Scale bar 100 μm. N=4 independent experiments. (C) Representative images of Diffquick stained cytospins of erythroid cells cells grown in liquid culture at the times indicated. Morphological analysis of erythroid cells shows the maturation stages during different days of the culture: Proerythroblast (red arrow), Basophilic erythroblast (blue arrow), Polychromatic erythroblast (green arrow), Orthochromatic erythroblast (orange arrow) & Mature cells (yellow arrow). The pictures were taken Leica DM6000 at 40x magnification, 20 μm scale bar. N= 4 independent experiments.

**Supplementary figure 3:**
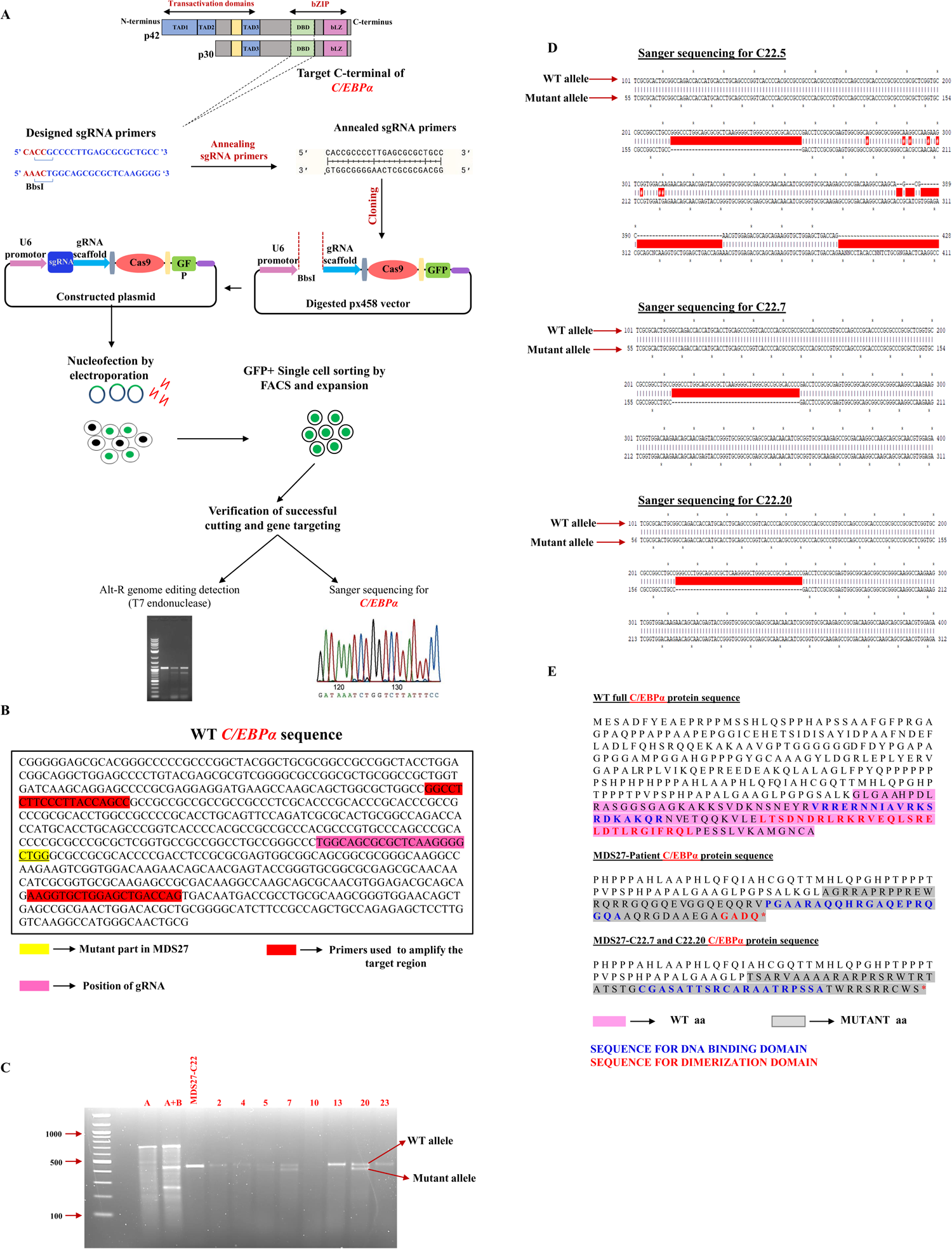
Generation of high-risk MDS27 isogenic clones by CRISPR-Cas9. (A) Schematic representation of engineering *C/EBPα* mid region mutation in MDS27-C22 using PX458 Cas9 plasmid. Guide RNA targeted mid region of C/EBPα were designed via an online tool (Trust Sanger Institute Editing database). Then, the sgRNA guide was cloned into an expression plasmid bearing both sgRNA scaffold backbone (BB) and Cas9, pSpCas9(BB) (PX458). The constructed plasmid was transfected into hiPSC MDS27-C22. T7I endonuclease assay and Sanger sequencing confirmed the generation of *C/EBPα* mutation. (B) Snapshot of C/EBPα sequence centre in the mutated region. Position of gRNA (pink), primers used to amplify the targeted region (red) and the deleted nucleotides (blue) after CRISPR-Cas9 are highlighted. (C) Mismatch cleavage assay T7EI was performed in different isolated clones. The Digestion reactions were analysed on an agarose gel. Sample 1 contains Control A homoduplexes PCR products, while Sample 2 contains homoduplexes and heteroduplexes of Control A and B PCR products. Expected DNA fragment size is indicated as WT, and that showing a deletion indicated as mutant allele. (D) Sanger sequencing of three clones (MDS27-C22.5, MDS27-C22.7 and MDS27-C22.20) with heterozygous mutation.

**Supplementary figure 4:**
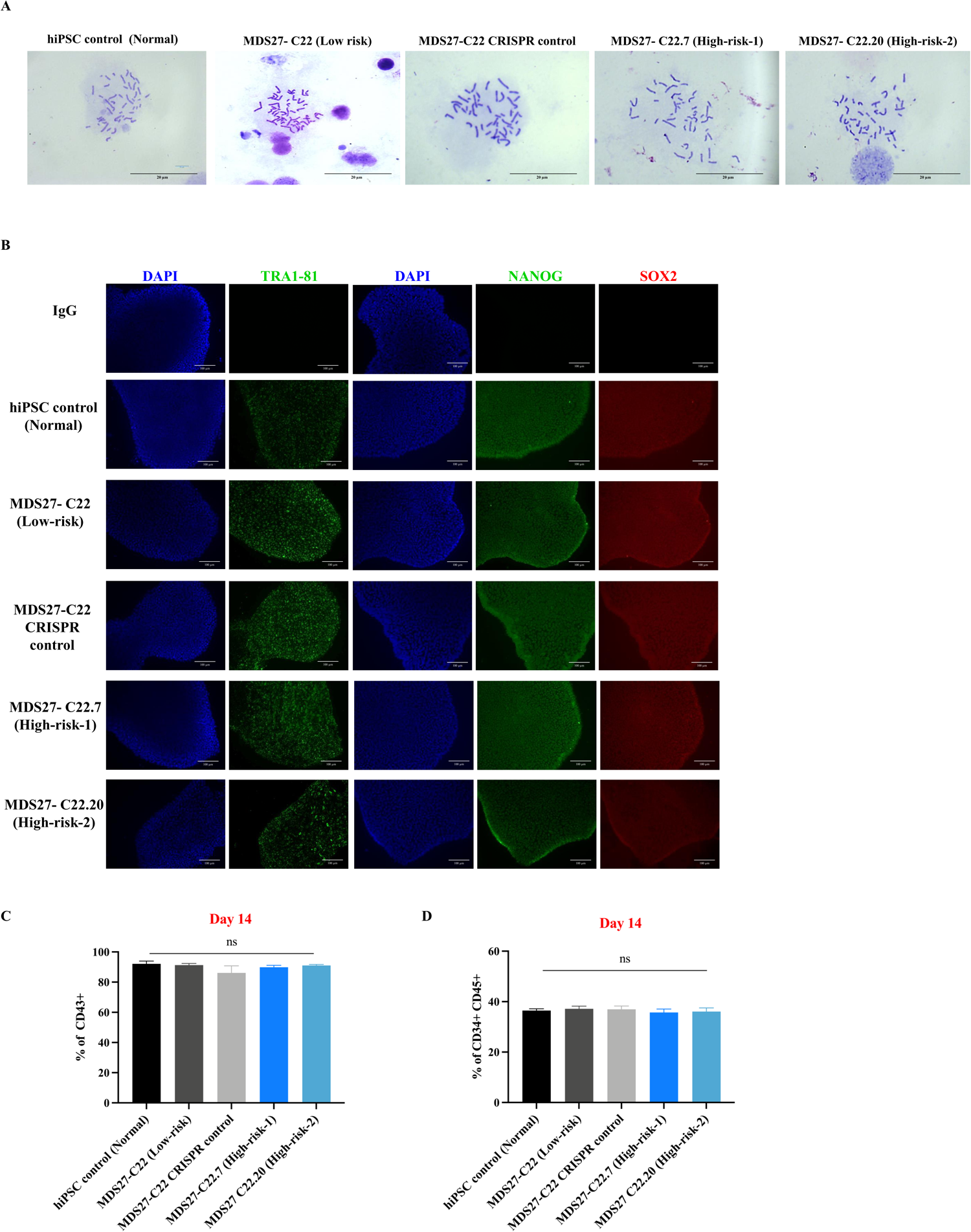
CRISPR-Cas9 system does not affect the chromosomal stability, the pluripotency of hiPSC and formation of HSPCs. (A) Chromosome spreads showing normal number of chromosomes for the indicated iPSCs lines. (>25 metaphases per sample per experiment), 20μm scale bar, 100x magnification, Leica DM6000 light microscope. (B) Representative immunofluorescence images after staining with pluripotency markers. DAPI staining is shown in blue. Scale bars, 100Cµm (n=3 independent experiments). (C) Bar graph showing the percentage of early hematopoietic population (CD43 ^+^) on day 14 of differentiation for the indicated iPSCs lines. Statistical results are presented as mean ± SEM. Ns= no significant, One-way ANOVA with multiple comparisons. N=3 independent experiments. (D) Bar graph showing the percentage of late hematopoietic population (CD34 ^+^ CD4 5^+^) on day 14 of differentiation for the indicated iPSCs lines. Statistical results are presented as mean ± SEM. ns= no significant, One-way ANOVA with multiple comparisons. N=3 independent experiments.

**Supplementary figure 5:**
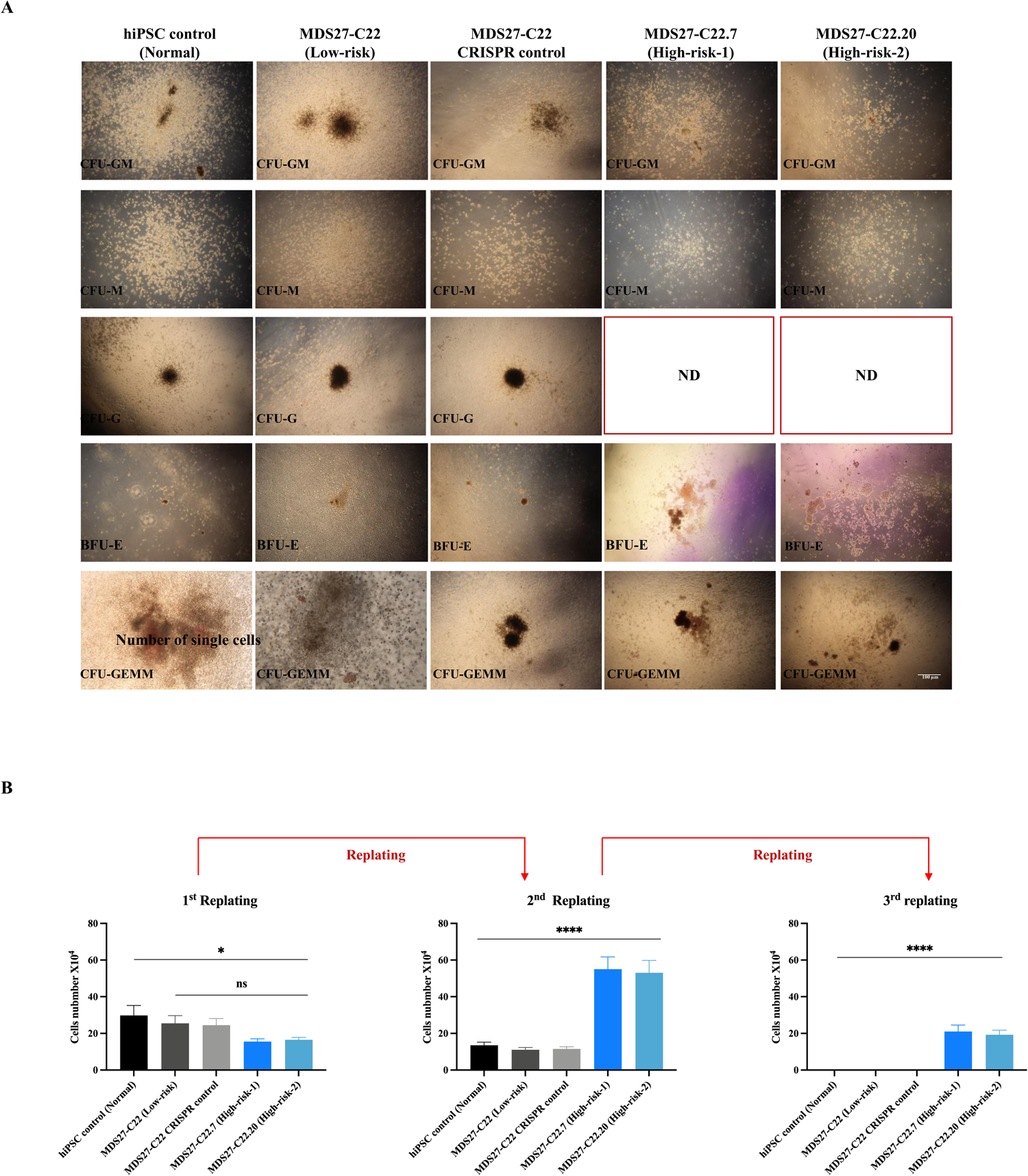
Further potential differentiation of HSPCs in semi-solid medium. (A) Representative images illustrating the morphology of CFUs scored after 14 days in semi-solid medium. scale bar 100 μm. n=4 independent experiments. (B) Number of the single cells that obtained aftere achreplating. Statistical represented as mean ± SEM. ns= no significant *p <0.05, ****p <0.0001, One-way ANOVA with multiple comparisons. N=3 independent experiments.

**Supplementary figure 6:**
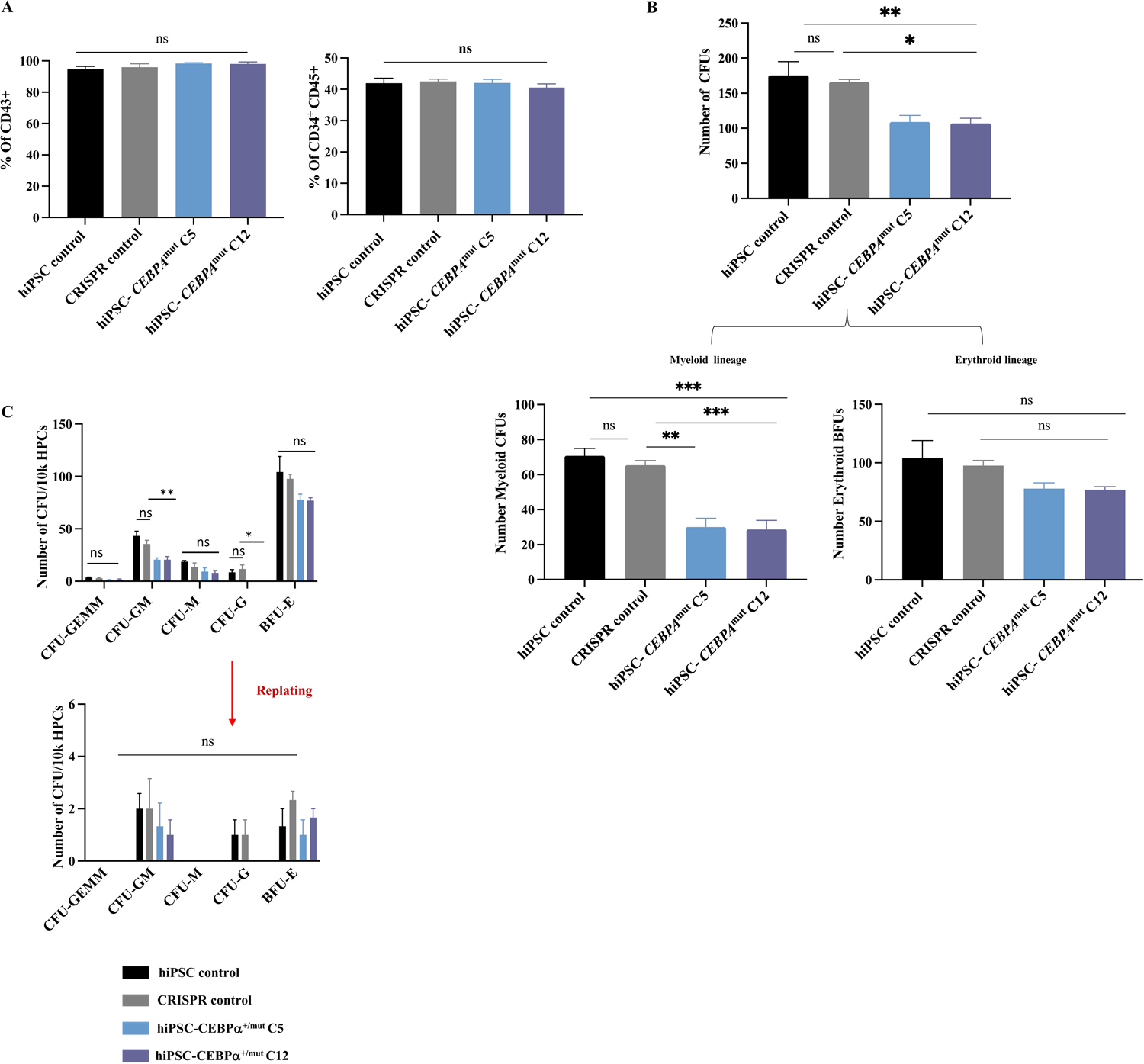
*C/EBPα* mutation does not confer self-renewal capacity to healthy HSPCs. (A) Bar graph showing the percentage of early hematopoietic population (CD43 ^+^) and late hematopoietic population (CD34 ^+^ C D 4^+^)5on day 14 of differentiation. Statistical results are presented as mean ± SEM. ns= no significant, One way AnOVA with multiple comparisons. N=3 independent experiments. (B) Total number of CFUs from 10 ^4^ –iPSC-HSPC cells grown for 14 days in semi solid medium. Statistical results are presented as mean ± SEM. * p< 0.05; **p< 0.01, t test. N= 3 independent experiments. (C) Number of CFUs obtained from healthy iPSC, healthy iPSC CRISPR control and healthy iPSC clones containing *C/EBPa^bZIP^* (C5 and C12) after first and second re-plating, each maintaining for 14 days. Mean and SEM are shown. **** p < 0.0001, ** p<0.001 and * p 0.05, t test. N=3 independent experiments.

**Supplementary Figure 7.**
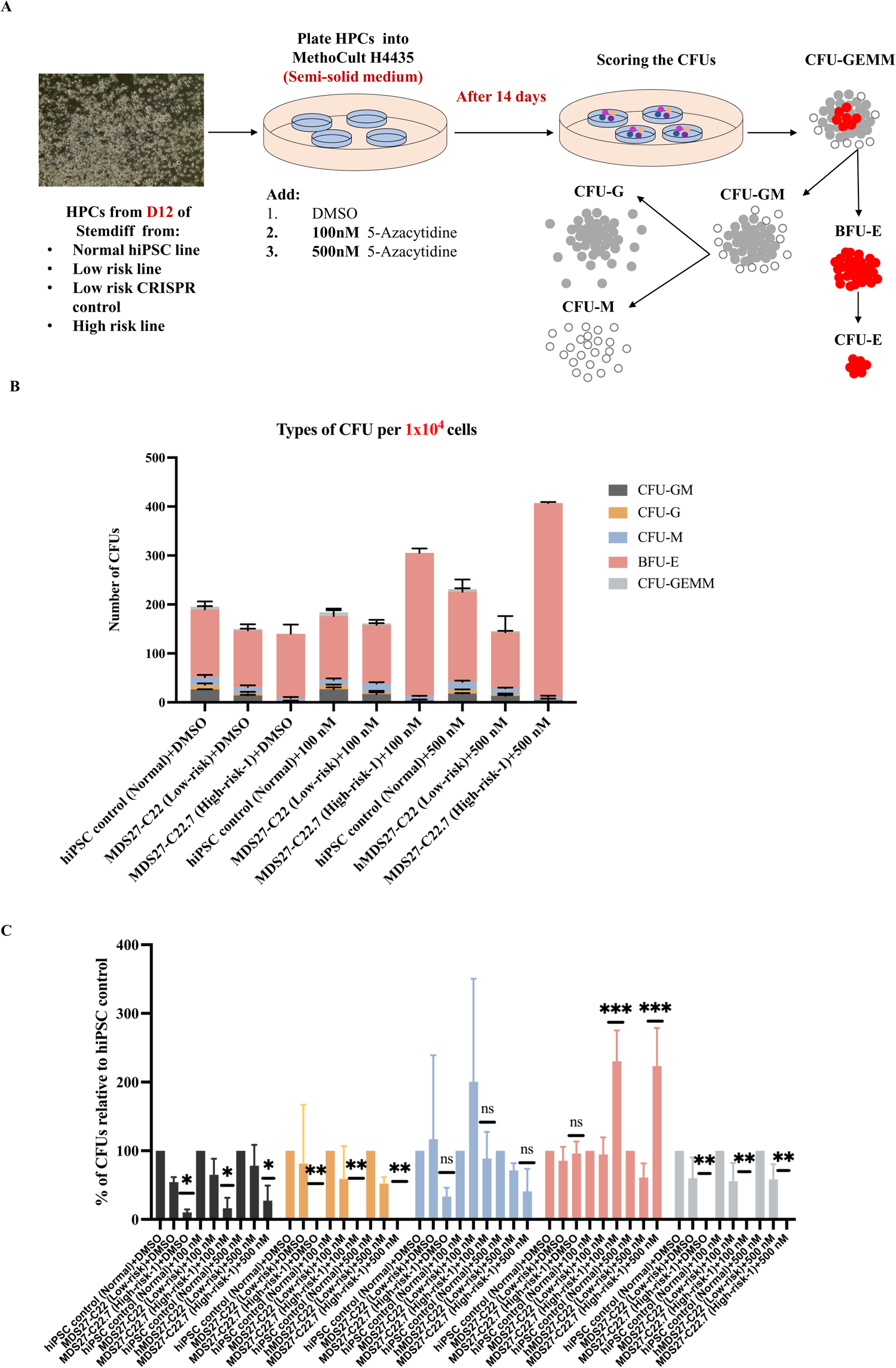
High-risk MDS-iPSC containing C/EBPα mutation are resistant to 5-Azacitine treatment. (A) Schematic representation of the experimental procedure for the colony forming assays. (B) Number of each type of CFUs after 14 days in semisolid medium. Mean and SEM of different lines are shown. N= 3 independent experiments. (C) Relative percentage of each type of CFUs for 1×10 ^4^ of HSPCs after 14 days in semisolid media. Statistical results are presented as mean ± SEM and **** p < 0. 0001, **p< 0.01, * p< 0.05 and (ns, no significant), Two-way ANOVA with multiple comparisons. N= 3 independent experiments.

**Supplementary figure 8.**
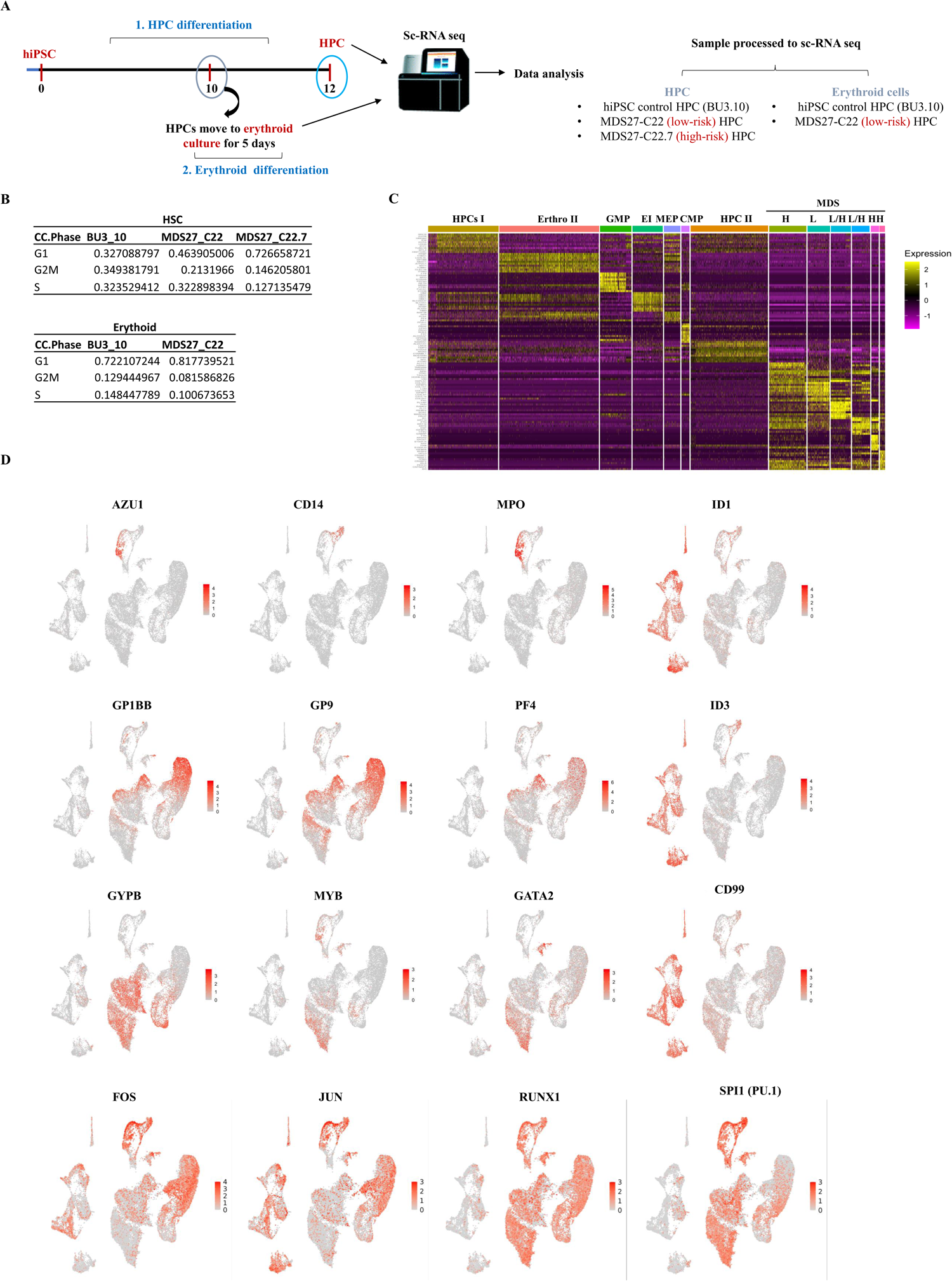
Distinct MDS transcriptome signature and clonal evolution during disease progression measured by single-cell transcriptome analysis. (A) Schematic representation of scRNAseq procedure. (B) Heatmap depicting the top 10 most upregulated marker genes identified in each scRNA-seq cluster. (C) Expression of indicated genes projected on the UMAP map. Colour intensity represents expression data log2 normalized unique molecular identifier (UMI) counts.

**Supplementary Figure 9:**
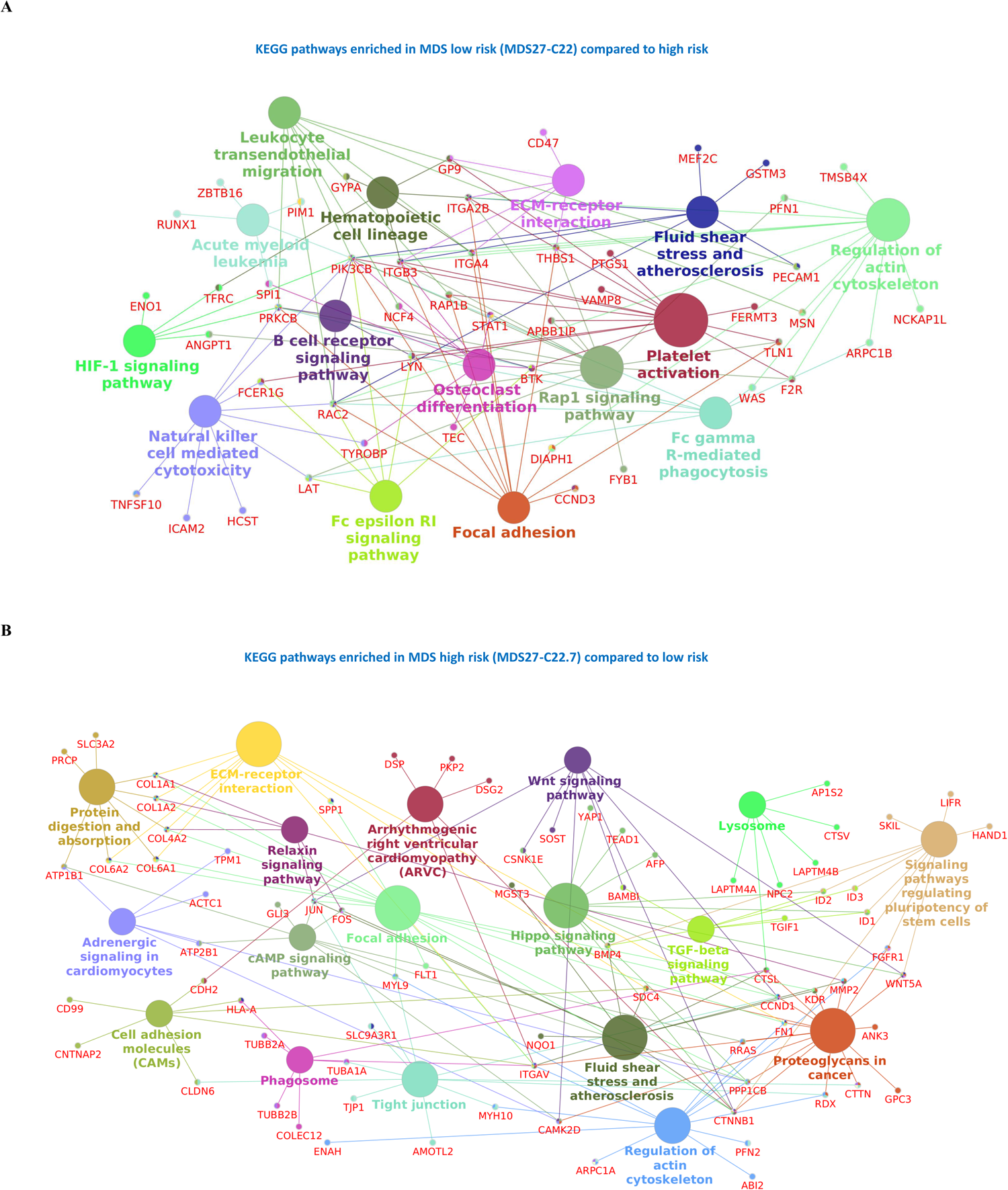
KEGG pathways for low risk and high risk HSPCs from Pseudo-bulk RNA-seq. A) KEGG pathway enrichment analysis of genes upregulated in Low risk MDS27-C22 iPSC compared to high risk MDS27-C22.7 iPSC. B) KEGG pathway enrichment analysis of genes upregulated in high risk MDS27-C22.7 compared to Low risk MDS27-C22 iPSC.

**Supplementary Table 1:** Genotyping of clones generated by Sendai virus and episomal reprograming Genotyping was performed by the Hospital La Fe, Valencia, Spain for generated clones from MDS27 peripheral blood sample on 2013, using an array of the 40 most mutated genes in AML. All clones harbour the same mutations as the original sample (2013).

**Supplementary Table 2:** Pseudo-bulk cluster analysis.

**Supplementary Table 3:** Myeloid signature for sc-RNAseq myeloid clusters.

